# NetworkExtinction: an R package to simulate extinction’s propagation and rewiring potential in ecological networks

**DOI:** 10.1101/2020.10.17.305391

**Authors:** M.Isidora Ávila-Thieme, Derek Corcoran, Erik Kusch, Simón P. Castillo, Fernanda S. Valdovinos, Sergio A. Navarrete, Pablo A. Marquet

## Abstract

1. Earth’s biosphere is currently undergoing drastic reorganisation as a consequence of the sixth mass extinction brought on by the Anthropocene. Impacts of local and regional extirpation of species have been demonstrated to propagate through the complex interaction networks they are part of, subsequently leading to secondary extinctions, exacerbating biodiversity loss. Contemporary ecological theory has developed several measures to analyse the structure and robustness of ecological networks under biodiversity loss. However, a toolbox for direct simulation and quantification of extinction cascades and the creation of novel interactions (i.e. rewiring) remains absent.
2. Here, we present *NetworkExtinction* - a novel R package which we have developed to explore the propagation of species extinctions sequences through ecological networks as well as quantify the effects of rewiring potential in response to primary species extinctions. With *NetworkExtinction* we have integrated ecological theory and computational simulations to develop functionality with which users may analyze and visualize the structure and robustness of ecological networks. The core functions introduced with *NetworkExtinction* focus on simulations of sequential primary extinctions and associated secondary extinctions while allowing for user-specified secondary extinction thresholds and realisation of rewiring potential.
3. With the package *NetworkExtinction,* users can estimate the robustness of ecological networks after performing species extinction routines based on several algorithms. Moreover, users can compare the number of simulated secondary extinctions against a null model of random extinctions. In-built visualizations enable graphing topological indices calculated by the deletion sequence functions after each simulation step. Finally, the user can define the degree distribution of the network by fitting different common distributions. Here, we illustrate the use of the package and its outputs by analyzing a Chilean coastal marine food web.
4. *NetworkExtinction* is a compact and easy-to-use R package with which users can quantify changes in ecological network structure in response to different patterns of species loss, thresholds, and rewiring potential. Therefore, this package is particularly useful to evaluate ecosystem responses to anthropogenic and environmental perturbations that produce non-random species extinctions.

## Introduction

Biological systems are commonly represented as complex networks of interactions (i.e., links between nodes representing species), through which matter and energy flow in a structured way, as in food webs (Benedek *et al.*, 2007; Pascual & Dunne, 2006; Proulx *et al.*, 2005) and mutualistic networks (González-Castro *et al.*, 2021; Schleuning *et al.*, 2016; Sebastián-González *et al.*, 2015). A myriad of perturbations, such as those produced by climate change and/or direct human activities, could lead in many cases to a local or global extinction of species or to severe reductions in abundance (Barnosky *et al.*, 2011; Costello *et al.*, 2016; May *et al.*, 1995; Pimm *et al.*, 2019; Scheffer *et al.*, 2001; Vitousek *et al.*, 1997). Such changes can deeply alter the energy fluxes at different temporal and spatial scales (Donohue *et al.*, 2016; Radchuk *et al.*, 2019; Venter *et al.*, 2016), modifying the ecological network components by adding or removing species and interactions, re-wiring, and changing interaction strengths. These impacts can be propagated through the ecological network and alter the stability and resilience of the ecosystem (Dunne *et al.*, 2002b).

Whether cascading effects are observed or not after removal or addition of a node depends, to some extent, on the complex structural attributes (also known as topological properties) that define the network (McWilliams *et al.*, 2019). Since species extinction and / or modification of their interactions may directly induce the degradation of ecosystem services, affecting human well-being, anticipating the potential propagation of these effects is of paramount importance (Barnosky *et al.*, 2011; Dirzo & Raven, 2003). Therefore, the understanding of ecosystem stability and resilience to different perturbations inducing species extinctions, has received considerable attention in the literature (Allesina & Pascual, 2009; Ávila-Thieme *et al.*, 2021; Curtsdotter *et al.*, 2011; Dunne *et al.*, 2002a; Hastings *et al.*, 2016; Jordan, 2009; Pimm *et al.*, 2019; Ramos-Jiliberto *et al.*, 2012; Roopnarine, 2006; Roopnarine *et al.*, 2007; Valdovinos, 2019; Valdovinos *et al.*, 2009).

### Topological Properties Shaping the Stability of Ecological Networks

The complexity of ecological networks imposes some challenges in developing an integrated framework and tools to study these systems. However, some general attributes that characterize most empirically constructed ecological networks do exist. For example, the impact of the number of species and connectance on ecological network robustness (measured as the number of secondary extinctions, see box 1 for definitions) have been emphasized by several authors but discrepancies still persist. While some studies suggest that increasing the number of species and connectance among them delay the onset of cascades of secondary extinctions (Dunne & Williams, 2009; Dunne *et al.*, 2002b; Estrada, 2007; Gilbert, 2009), others show the opposite relationship (Pires *et al.*, 2015; Sauve *et al.*, 2014; Staniczenko *et al.*, 2010). Thébault & Fontaine (2010) propose that these discrepancies may be driven by the type of network (e.g. trophic versus mutualistic networks), which necessitates a different treatment of mutualistic and trophic networks when studying extinction cascades.

##### Box 1: Definitions relevant to the *NetworkExtinction* R package workflows

- **Network Robustness** - A measure of the maintenance of network structure in the face of perturbations and quantified here as the number of species (nodes) lost as a consequence of primary species extinctions.
- **Interaction Type** - a link between two nodes reflecting the type of relationship involved. *NetworkExtinction* handles mutualistic (+/+) and trophic / parasitic (-/+) interaction types. For a more exhaustive overview of interaction types, consult Morales-Castilla *et al.* (2015).
- **Interaction strength** - The direct effect that nodes have on each other’s demography (Morales-Castilla *et al.*, 2015), fitness (de Santiago-Hernández *et al.*, 2019), or resource acquisition/transfer of energy (Heymans *et al.*, 2016). *NetworkExtinction* implicitly treats interaction strength as the effect that nodes have on each other’s persistence.
- **Extinction Threshold** - *NetworkExtinction* treats an extinction threshold as a percentage of interaction strength loss (relative to the total interaction strength at onset of extinction simulation) which a vertex may loose before becoming secondarily extinct (Schleuning *et al.*, 2016).
- **Rewiring capability** - Rewiring is the process by which a vertex may allocate interaction strength linked to a link which is removed due to a loss of interaction partner to an entirely new or already linked partner thereby increasing the interaction strength of new or already existing links in a network (Fründ, 2021; Schleuning *et al.*, 2016; Staniczenko *et al.*, 2010).

Similarly, theoretical models show that the degree distribution (i.e., distribution of links per node) of ecological networks is strongly associated with their robustness to species loss (Sole & Montoya, 2001). Usually, degree distributions follow a fat-tailed distribution (Bascompte, 2009; Dunne *et al.*, 2008). However, power-law degree distributions where super-connected nodes are more common, are more vulnerable to the removal of the most connected nodes (de Santana *et al.*, 2013; Dunne *et al.*, 2002a; Estrada, 2007; Sole & Montoya, 2001). More generally, a directed attack to the nodes with higher degree can have larger whole-scale consequences in the network (Albert & Barabási, 2002; Albert *et al.*, 2000). Identifying the best model to describe an empirical degree distribution has been an active research area and is a common task in the analysis of ecological networks.

### The Need for Simulation Approaches in Extinction Analyses

While the assessment of ecosystem robustness through topological metrics of ecological networks is computationally inexpensive (there are packages that evaluate it easily, see Table S1), relying on topological metrics alone may be misleading considering the differ implications given network types (Thébault & Fontaine, 2010), and the sometimes weak connection between these metrics and real (empirical) network resiliency to extinctions (e.g. Ávila-Thieme *et al.* 2021). Alternatively, assessments of the importance of species on ecological network persistence can be carried out by simulating a sequence of species removal and evaluating the consequences on network topology and robustness. Such extinction simulations are computationally much more expensive than the single-step calculation of topological metrics, but capable of rendering a more direct quantification of network robustness. Simulating responses to an extinction scenario involves a number of computationally demanding steps, which can be challenging and time consuming for researchers. Some node metrics (e.g., node degree) change dynamically throughout the sequential extinction simulation and need to be recalculated after each simulation iteration.

Several indices and open source R packages have been created to visualize and analyze the topology and dynamics that occur within the networks (Table S1). However, despite the recent diversification of analytical tools, we still lack a way to asses ecological network robustness to a ‘user-defined’ sequence of species loss implemented in an open source platform, such as R, using approaches that integrate key elements for ecological networks, such as the type of interactions, extinction thresholds and network rewiring. To fill this software and functionality gap, we have developed an open-source R package that facilitates the exploration of ecological network (trophic and mutualistic) robustness (see box 1 for definitions) and changes in attributes following the removal or extinction of nodes in complex ecological networks. Here, we present the *NetworkExtinction* R package, which quantifies changes in ecological networks both topologically and returns post-extinction networks according to simulated extinction sequences and their consequences.

### Interaction Types, Extinction Thresholds & Network Rewiring

Ecological network types are manifold and may be classified the interaction type they encode (e.g., trophic or mutualistic), how many levels of organisms they represent (e.g., bipartite or multilayer networks), whether they quantify interactions or simply denote their presence/absence (i.e., weighted vs. binary networks), and whether they represent realized or potential interactions. To best represent network changes in response to node removal, co-extinction simulation frameworks ought to account for the network-type specific changes in network cascade responses.

When considering simulations of extinction cascades, the core use of the *NetworkExtinction* package, it is thus critical to focus on three important aspects of networks (see box 1 for definitions): (1) interaction types, (2) interaction strength introducing extinction thresholds, and (3) potential rewiring of lost interactions enabling continued persistence of species.

Interaction Types are the cornerstone of most ecological network research as they greatly impact how links between nodes are interpreted biologically and subsequently impact the consequences of loss of connections in extinction cascades. For example, in trophic networks, basal species may lose all associated predators, resulting in isolated nodes, but not in their extinction. In a mutualistic network, on the other hand, loss of all connections will inevitably lead to extinction of any node (given that the network encodes interactions required for survival) (Carpentier *et al.*, 2021; Schleuning *et al.*, 2016).

However, a species does not necessarily have to lose all its interaction partners to be in danger of going extinct (Bascompte & Jordano, 2007). Such extinction thresholds may exist either globally for all nodes within a network or individually for each node separately. For example, a predator species may lose all but its main prey species and still continue to thrive, but die out when losing access to its main prey. In this case, an extinction threshold ought to incorporate interaction strengths (i.e, link weights in network representation) which will indicate which interaction partner is most important for the target node.

Contrary to the discussion of extinction consequences so far, there is also potential for novel interactions or changes in established interaction strengths, which may manifest as the rewiring of networks in response to primary extinctions (Bartley *et al.*, 2019; Ramos-Jiliberto *et al.*, 2012; Staniczenko *et al.*, 2010; Strona & Bradshaw, 2018; Valdovinos, 2019; Vizentin-Bugoni *et al.*, 2020). Rewiring potential has recently received increased attention from the ecological network community as a possible mechanism by which the impacts of the Anthropocene may be abated. At its core, rewiring of interactions is a process by which links that are lost due to removal of a node may be reallocated either to novel interaction partners or combined with existing interactions. Recalling the previous example of a predator losing access to its main prey item, when considering rewiring potential, this predator may shift to preying on other prey which is already contained in its diet, or interact with entirely new prey instead of going extinct.

Most contemporary analyses of ecological networks and simulations of extinction consequences incorporate one or two of these considerations (interaction type, extinction threshold, and rewiring), but rarely all three (Schleuning *et al.*, 2016). We suggest that this is a consequence of the complexity of identifying appropriate thresholds of extinction risks and rewiring potential that can be realised as well as complexity of analysis tools required to incorporate these mechanisms. To our knowledge, the *NetworkExtinction* package is the first implementation of all these considerations into one easy- and free-to-use software package.

### The *NetworkExtinction* R Package

The *NetworkExtinction* package analyzes ecological networks representing species as nodes and their interactions as links. The links within the networks can be weighted or binary. Using this input (formatted either as an adjacency matrix or a network object), the *NetworkExtinction* package simulates species extinctions sequences (`SimulateExtinctions` and `RandomExtinctions` functions). Non-random extinctions can be simulated either as a static (“`Ordered`” method) or flexible (“`Mostconnected`” method) process. In doing so, the *NetworkExtinction* package interacts with other R packages, especially with the *network* package (Butts *et al.*, 2008). *NetworkExtinction* also visualizes the results (`ExtinctionPlot` function) and compares them between the different methods (`CompareExtinctions` function). Finally, *NetworkExtinction* fits the network degree distribution (`DegreeDistribution` function). See Figure 1 for a visual representation of this functionality.

**Figure 1:**
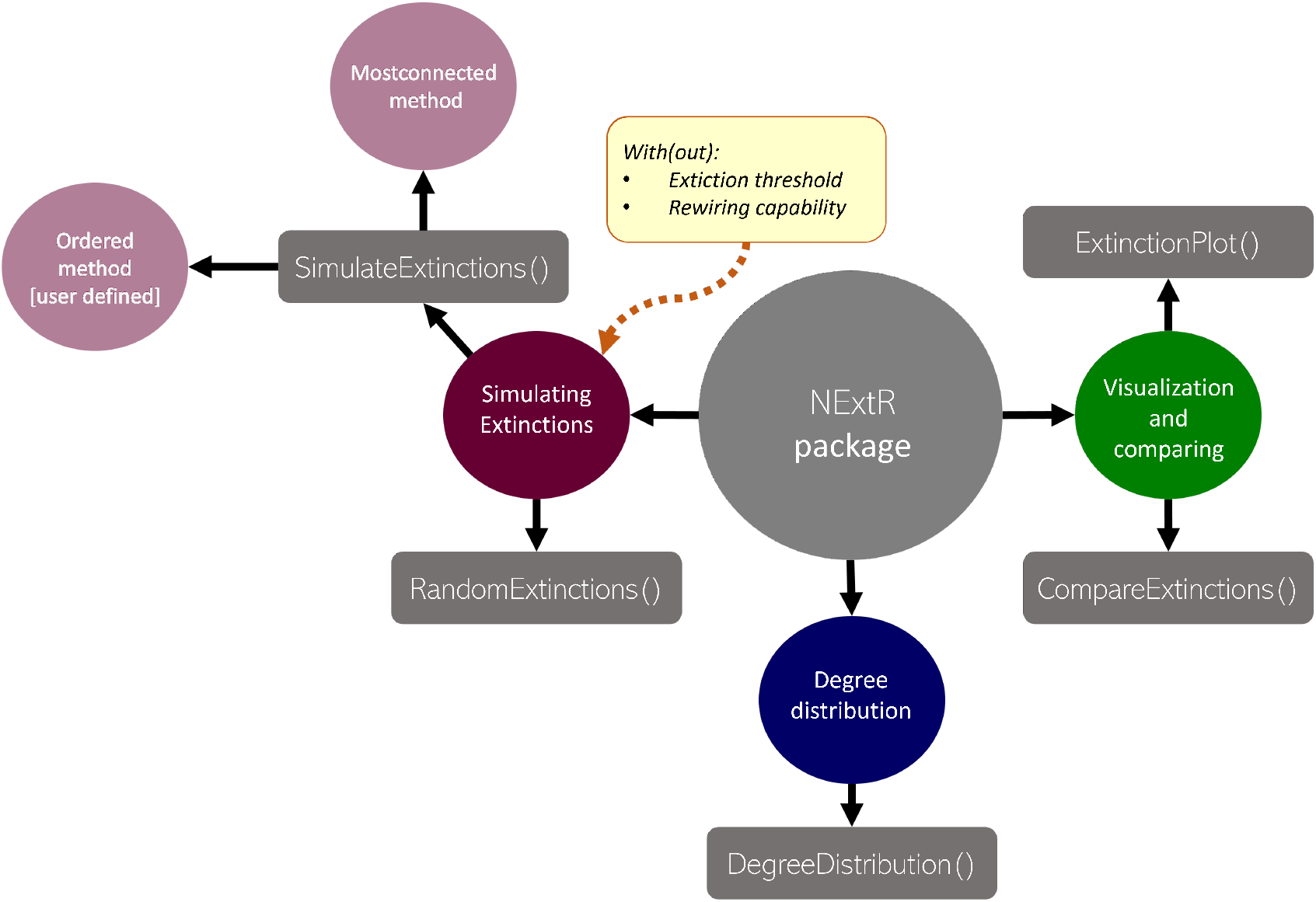
Synthesis and functions of the *NetworkExtinction* package and its functions.

When executing a simulation of extinction cascades using the *NetworkExtinction* package, users can specify (1) what interaction type (i.e., trophic or mutualistic) is being analysed, (2) whether to consider a species extinction threshold and (3) whether to simulate link rewiring. In the case of trophic ecological networks, only bottom-up trophic cascades (Berg *et al.*, 2015; Curtsdotter *et al.*, 2011; Dunne *et al.*, 2002b) are modelled (i.e., loosing predator species does not affect the survival of a prey node although it may become disconnected from the network).

Here, we demonstrate the functionality and outputs of the *NetworkExtinction* package using an empirical marine intertidal rocky shore trophic network (hereafter, “chilean_intertidal”), which contains 107 species forming 1381 realised trophic interactions (Ávila-Thieme *et al.*, 2021; Kéfi *et al.*, 2015). For a use-case of mutualistic network analyses with the *NetworkExtinction* package, see Kusch & Ordonez (2022). In the following, we focus on the implementation of a basic workflow with the *NetworkExtinction* package and how to augment extinction simulations with consideration of extinction thresholds and rewiring mechanisms. For a detailed overview of the functions within the R package, their input, and output, please refer to the supplementary material (section “Package Functions and Arguments”).

### The Basic Workflow

The NetworkExtinction package is hosted on CRAN and can be installed and loaded thusly:

~~~
R> install.packages(“NetworkExtinction”)
R> library(NetworkExtinction)
~~~

#### Extinction Functions

Two of the five functions contained in the *NetworkExtinction* package are used to simulate extinction cascades and measure ecological network topology and robustness after simulating a given species deletion sequence corresponding to primary extinctions and identifying secondary extinctions. These functions are called `SimulateExtinctions` and `RandomExtinctions`.

##### The SimulateExtinctions() Function

`SimulateExtinctions` enables the user to remove nodes from the network based on the following two deletion sequences: 1) species’ degree (“`Mostconnected`” method), 2) a user-defined order (“`Ordered`” method).

###### Mostconnected Extinction Order

Using the “`Mostconnected`” method, the `SimulateExtinctions` function first identifies the most connected species via the degree of its corresponding node (i.e., number of links attached to the node). This node is then removed from the network, and the function checks whether other species are now going extinct according to user specifications of the function (having become completely unconnected, in the default case shown here). This step is repeated until the entire network is unconnected. At each step, `SimulateExtinctions` recalculates node-degree for each extant species to re-identify the next most connected node up for primary removal (see code chunk 2 in the supplementary material).

The `SimulateExtinctions` function returns two objects: (1) a data frame (…`$sims`) containing topological metrics of the network after every step of species removal (Table 1) as well as (2) the reduced network (…`$Network`) corresponding to the portion of the original network extant after removal of primarily and secondarily extinct species. The “`Mostconnected`” method of `SimulateExtinctions` for the Chilean intertidal food web results in complete network annihilation after primary removal of the 37 most connected species. Consequently, the reduced network is empty.

**Table 1:**
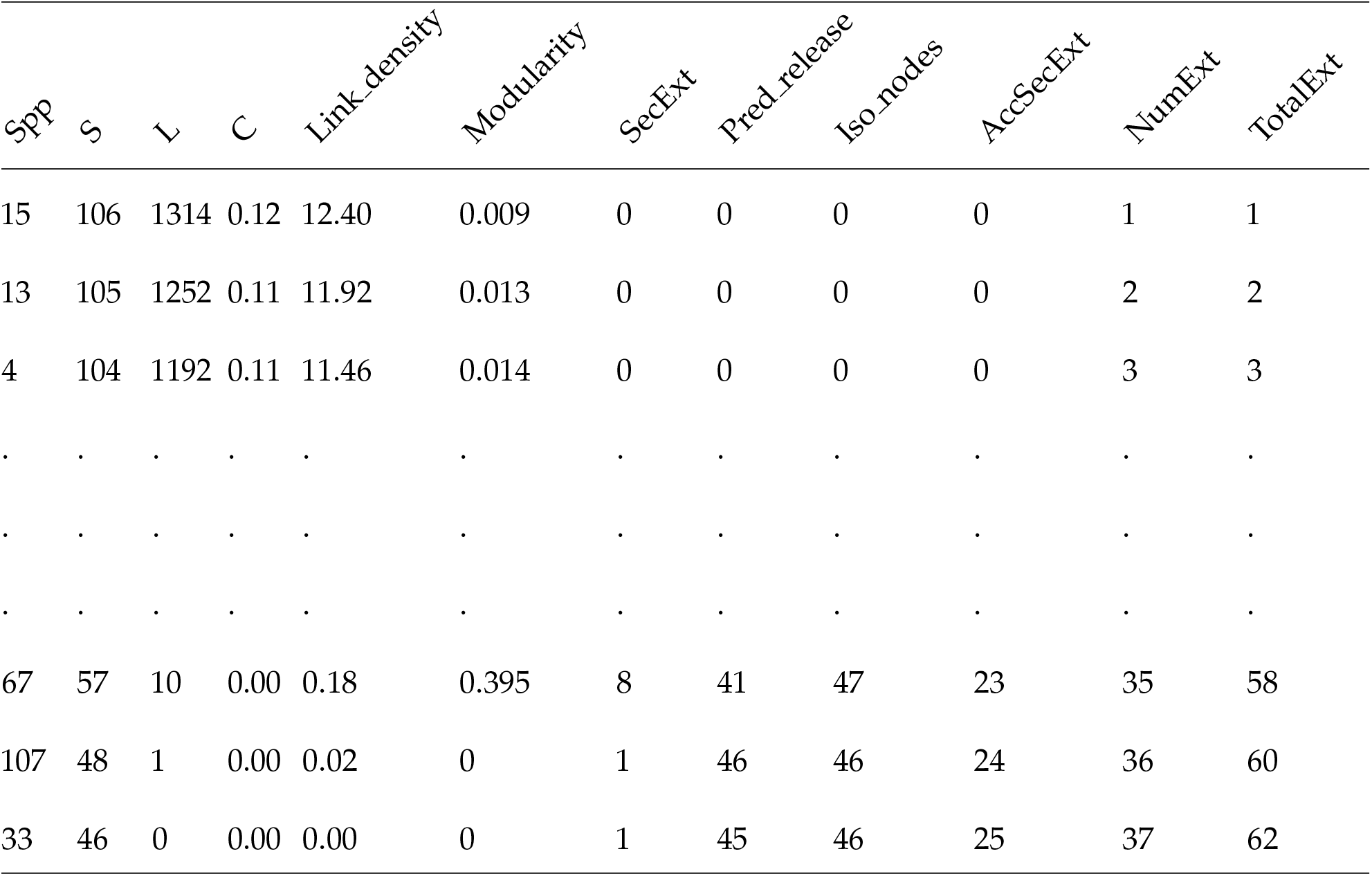
Summarised results of the *SimulateExtinctions* function with the *“Mostconnected”* method for the intertidal food web, showing the first and last three rows of the original data frame (see full results in Table S2). Spp: node position of the extinct species, S: richness, L: number of links, C: connectance, Link_density: link density (L/S), SecExt: secondary extinctions, Pred_release: predation release, Iso_nodes: isolated nodes, AccSecExt: cumulative number of secondary extinctions, NumExt: cumulative number of primary extinctions, TotalExt: number of total extinctions. See full results in Table S2 and code to produce this output in code chunk 2 in the supplementary material.

| Spp | S | L | C | Link_density | Modularity | SecExt | Pred_release | Iso_nodes | AccSecExt | NumExt | TotalExt |
| --- | --- | --- | --- | --- | --- | --- | --- | --- | --- | --- | --- |
| 15 | 106 | 1314 | 0.12 | 12.40 | 0.009 | 0 | 0 | 0 | 0 | 1 | 1 |
| 13 | 105 | 1252 | 0.11 | 11.92 | 0.013 | 0 | 0 | 0 | 0 | 2 | 2 |
| 4 | 104 | 1192 | 0.11 | 11.46 | 0.014 | 0 | 0 | 0 | 0 | 3 | 3 |
| . | . | . | . | . | . | . | . | . | . | . | . |
| . | . | . | . | . | . | . | . | . | . | . | . |
| . | . | . | . | . | . | . | . | . | . | . | . |
| 67 | 57 | 10 | 0.00 | 0.18 | 0.395 | 8 | 41 | 47 | 23 | 35 | 58 |
| 107 | 48 | 1 | 0.00 | 0.02 | 0 | 1 | 46 | 46 | 24 | 36 | 60 |
| 33 | 46 | 0 | 0.00 | 0.00 | 0 | 1 | 45 | 46 | 25 | 37 | 62 |

###### User-Defined Extinction Order

Supplying a user-defined order to `SimulateExtinctions` is particularly useful when knowledge about extinction risks of species exists or is inferred from species’ traits (e.g. size, trophic position). In contrast to the “`Mostconnected`” method of the `SimulateExtinctions` function, the “Ordered” method does not change the initial extinction order, but treats it as static. Here, we supply the 60 most connected species who aren’t top predators in the Chilean intertidal network (see code chunk 3 in the supplementary material and Table S3 for the full extinction sequence).

Regardless of the selected method, the `SimulateExtinctions` function returns the same kind of output previously described. However, having supplied a primary extinction order that does not include all nodes in the original network and whose extinction simulation did not lead to total network annihilation, we can also assess the post-extinction simulation network (Figure 2).

**Figure 2:**
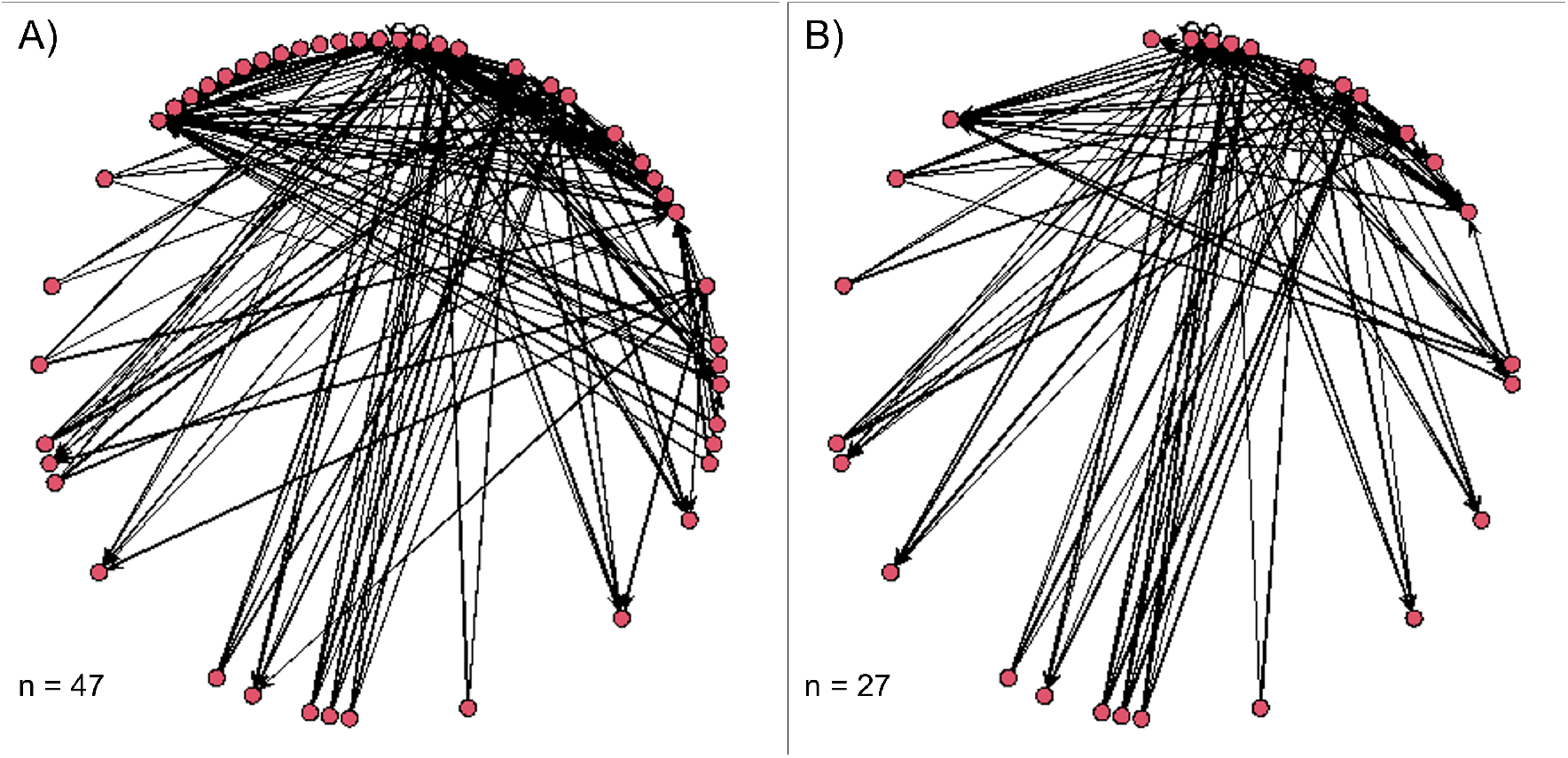
Post-extinction networks representative of removal of the 60 most connected non-top predator species from the Chilean intertidal network. A) Reduced network following removal of only primary extinction nodes. B) Reduced network obtained via the *SimulateExtinctions* function which also accounts for secondary extinctions. n = number of resulting nodes. See code chunk 4 in the supplementary material for generation of these networks and plots.

##### Random Extinctions

The second extinction simulation function - `RandomExtinctions` - allows users to simulate the removal of a number of nodes based on a random deletion sequence. The output of this function is particularly useful for establishing effect sizes of non-random deletion sequences (see code chunk 5 in the supplementary material).

The function returns a data frame (Table 2) and a `plot` (when the optional plot argument is set to `TRUE` (see code arguments in supplementary material) with the mean of secondary extinctions for each removal step averaged through all the simulations.

**Table 2:**
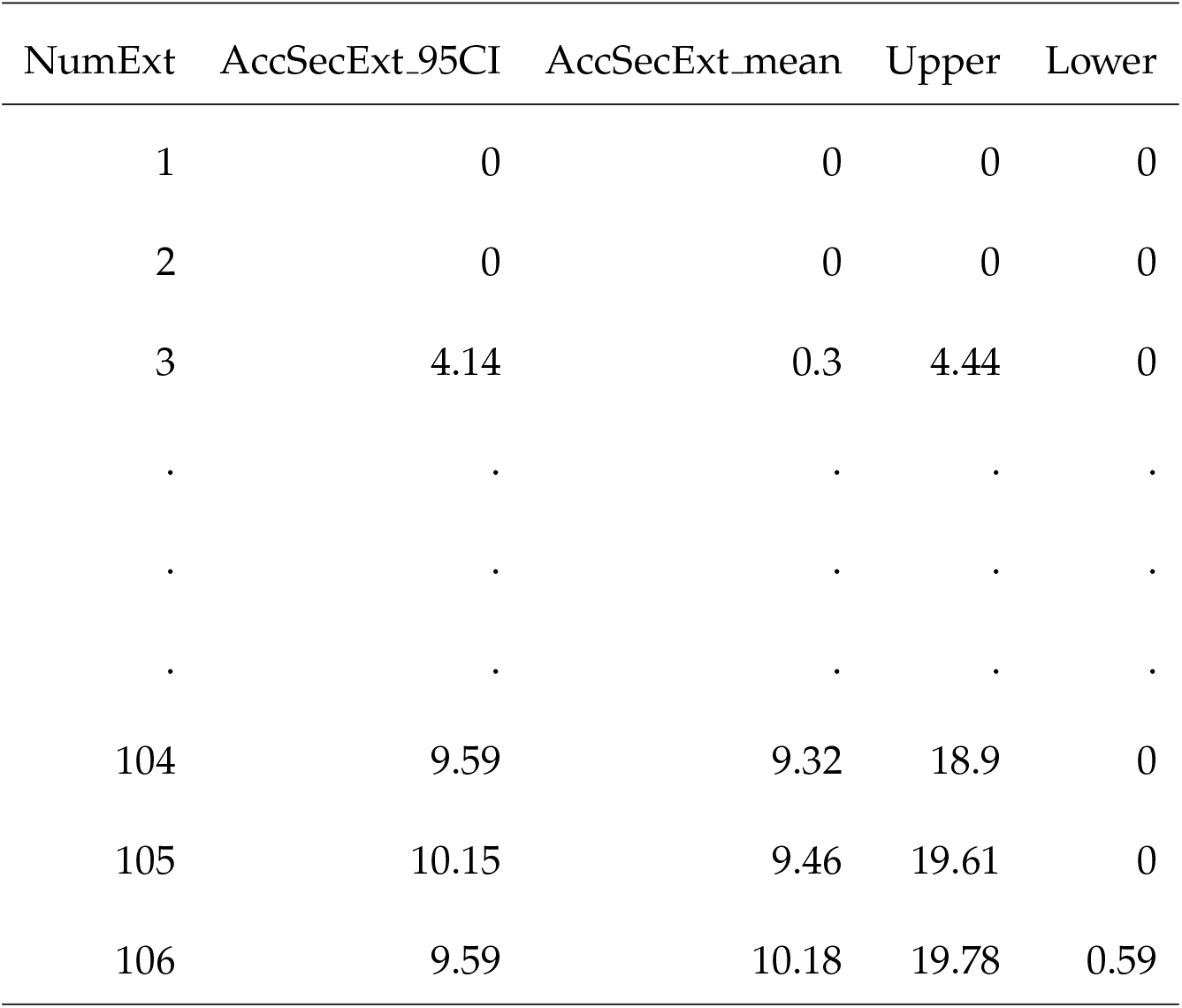
Summarised results of the *RandomExtinction* function for the intertidal food web, showing the first and last three rows. NumExt: cumulative number of primary extinctions, AccSecExt_95CI: cumulative 95% of confidence intervals of the secondary extinctions among all the simulations performed, AccSecExt_mean: cumulative average of secondary extinctions among all the simulations performed, Upper & Lower: lower and upper limit of the [mean + 95% CI], respectively. See the full results in Table S4 and the code to produce this output in code chunk 5 in the supplementary material.

#### Analysis & Visualization Functions

Two more functions contained in the *NetworkExtinction* package are used to visualize and analyze ecological networks and their extinction sequences beyond simulations of extinction cascades. These are called `ExtinctionPlot` and `CompareExtinctions`.

##### The ExtinctionPlot() Function

The `ExtinctionPlot` function is particularly useful for visualizations of extinction simulation outcomes as obtained through `SimulateExtinctions`. Using this function, users can plot any of the topological metrics that `SimulateExtinctions` calculates at each simulation step against the progress of the extinction simulation along the extinction order (see code chunk 6 in the supplementary material). As such, this function can visualize all columns displayed in the standard `SimulateExtinctions` output (Table 1). As an example, we plot the link density of the intertidal food web at each removal step using the “`Mostconnected`” deletion sequence of the `SimulateExtinctions` function (Figure 3).

**Figure 3:**
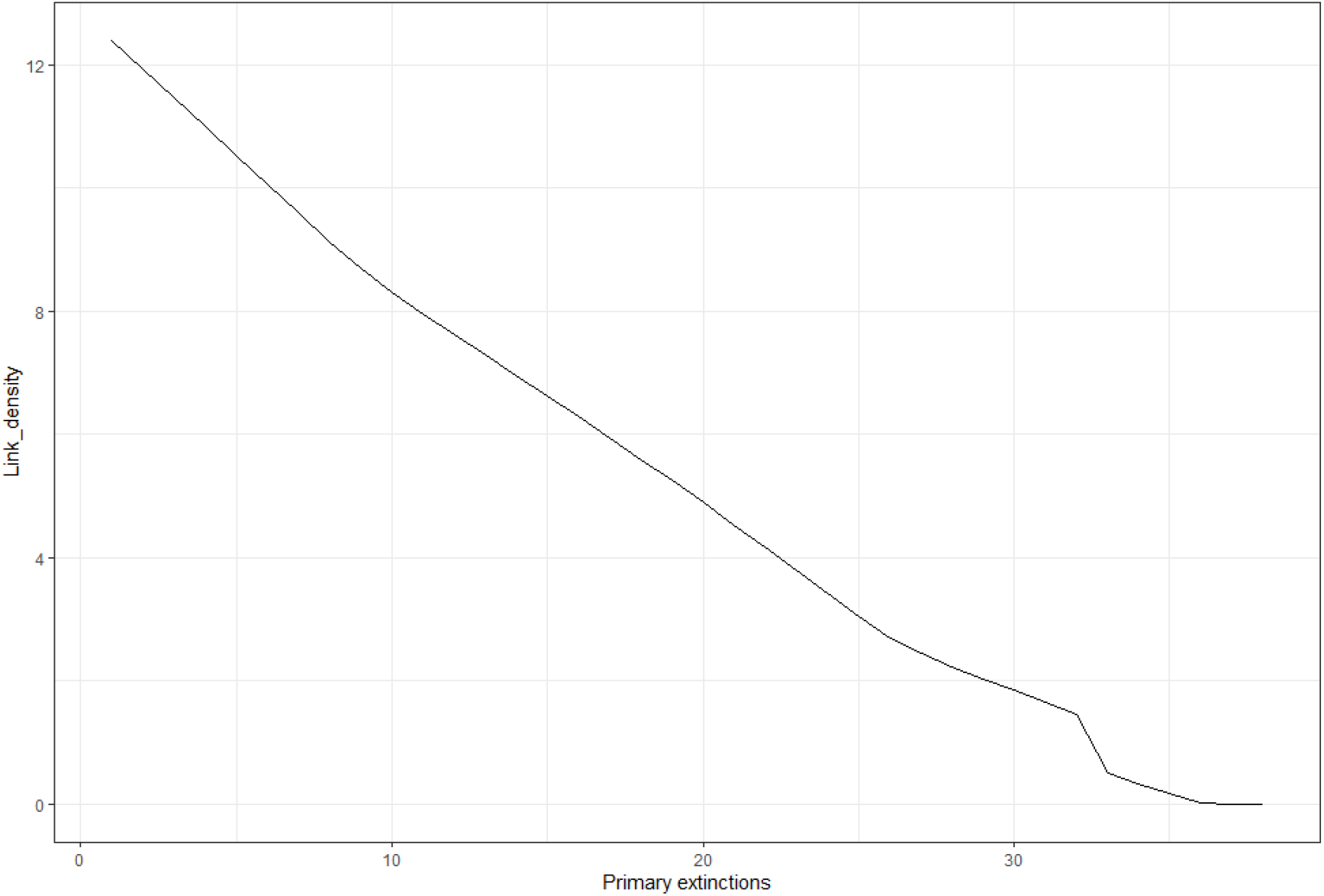
Links density after each removal step (primary extinctions) in the intertidal food web using the *“Mostconnected”* method of the *SimulateExtinctions* function and visualized using the *ExtinctionPlot* function.

##### The CompareExtinctions() Function

The `CompareExtinctions` function compares the number of secondary extinctions produced by either of the two options of the `SimulateExtinctions` function, against a set of random deletion sequences (see code chunk 6 in the supplementary material). This comparison is returned as a figure (Figure 4). Here, we compare the secondary extinctions produced by the random deletion sequences (`RandomExtinctions`) with the extinctions produced by the “`Mostconnected`” deletion sequence of the `SimulateExtinctions` function. In this example, Figure 4 shows clearly that primary extinction of the most connected species has a more drastic effect on the rate of secondary extinction accumulation than would be expected following random primary extinctions.

**Figure 4:**
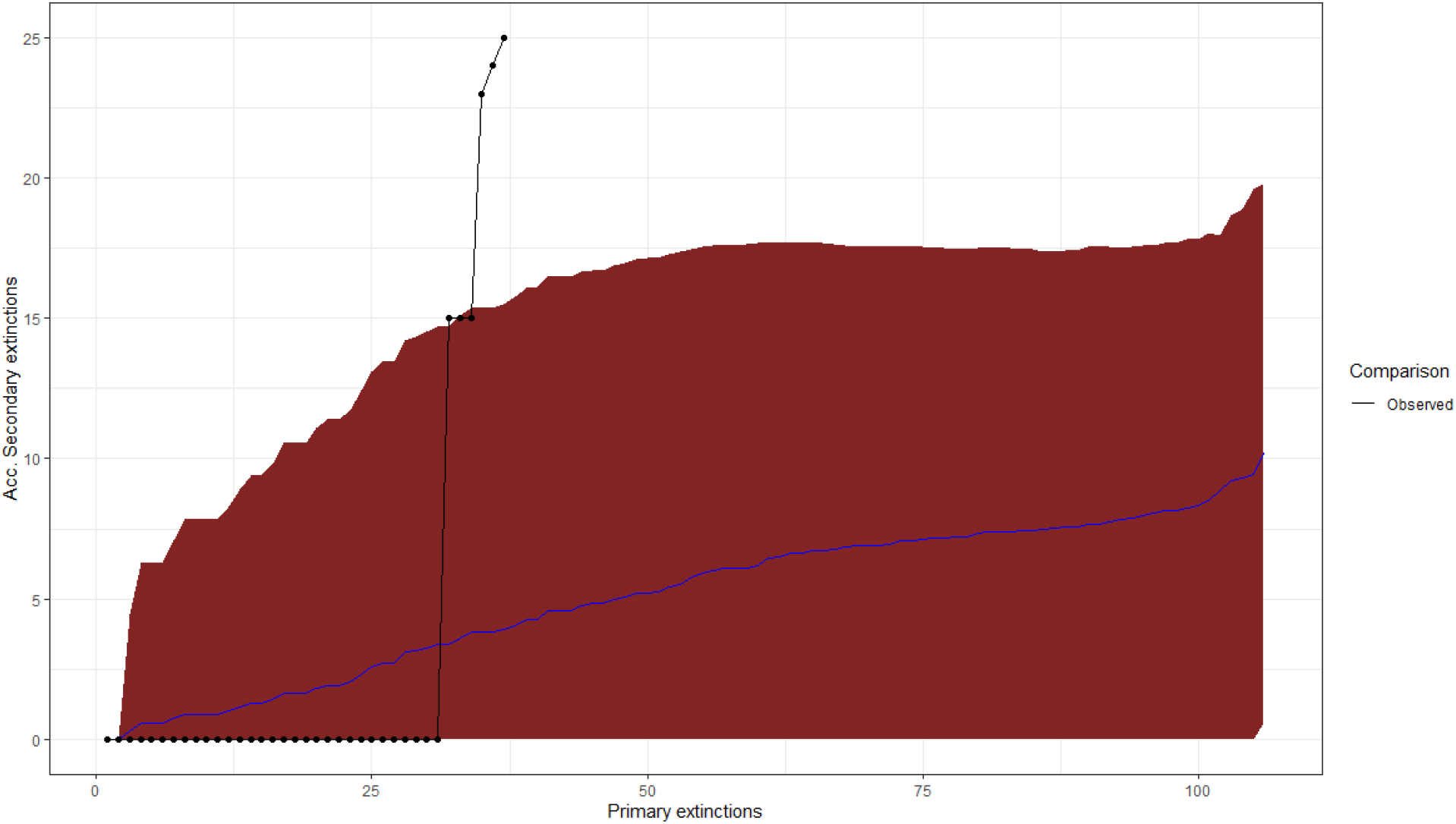
Comparison of the cumulative secondary extinctions after each removal step (Primary extinctions, defined by user) between the random (Null hypothesis) and *“Mostconnected”* (Observed) deletion sequence in the intertidal food web using the *CompareExtinctions* function. The blue line is the average (±95%CI [red area]) of secondary extinctions of the null model and the black line following the dots represents the secondary extinctions of the observed model.

#### Degree Distribution

The final function contained in the *NetworkExtinction* package - `DegreeDistribution` - fits the degree distribution of the network using two approaches: linear (on log-transformed data) and non-linear regression (see code chunk 7 in the supplementary material).

Different statistical approaches have been proposed to fit the degree distribution, such as maximum likelihood (Clauset *et al.*, 2009), ordinary least squares, or linear versus non-linear regression (Xiao *et al.*, 2011). As in other fields, the use of linear and non-linear regressions has been controversial (Xiao *et al.*, 2011). Some have suggested that the linearization using a logarithmic scale is flawed and that instead, the analysis should be conducted on the original scale using non-linear regression methods (Xiao *et al.*, 2011). In part, this is because when using linear regressions (LR) on log-transformed data the error distribution may not meet the assumptions needed to statistically compare across different models; hence, a second group of approaches considers the use of non-linear regression using general least squares, in combination with Akaike’s information criteria to select the best model that fits the degree distribution.

`DegreeDistribution` incorporates these considerations in its three data frames outputs (`models, params`, and `DDvalues`) with:

- `models`: Comparison of the AIC and normal distribution of the residual assumption test between the different distributions tested (Table 3).
- `params`: The statistical parameters of each model (Table 4) corresponding to *P*_(*k*)_ = *ck^β^* (non-linear power-law models), log *P*(*k*) = *β* log *k* + *c* (linear power-law models) and *P*_(*k*)_ = *e*^λ*k*+*c*^ (non-linear exponential distribution models), log *P*_(*k*)_ = λ*k* + *c* (linear exponential distribution models).
- `DDvalues`: The degree distribution with the observed values and the value of each fitted model (visualised automatically by the function as seen in Figure 5).

**Table 3:**
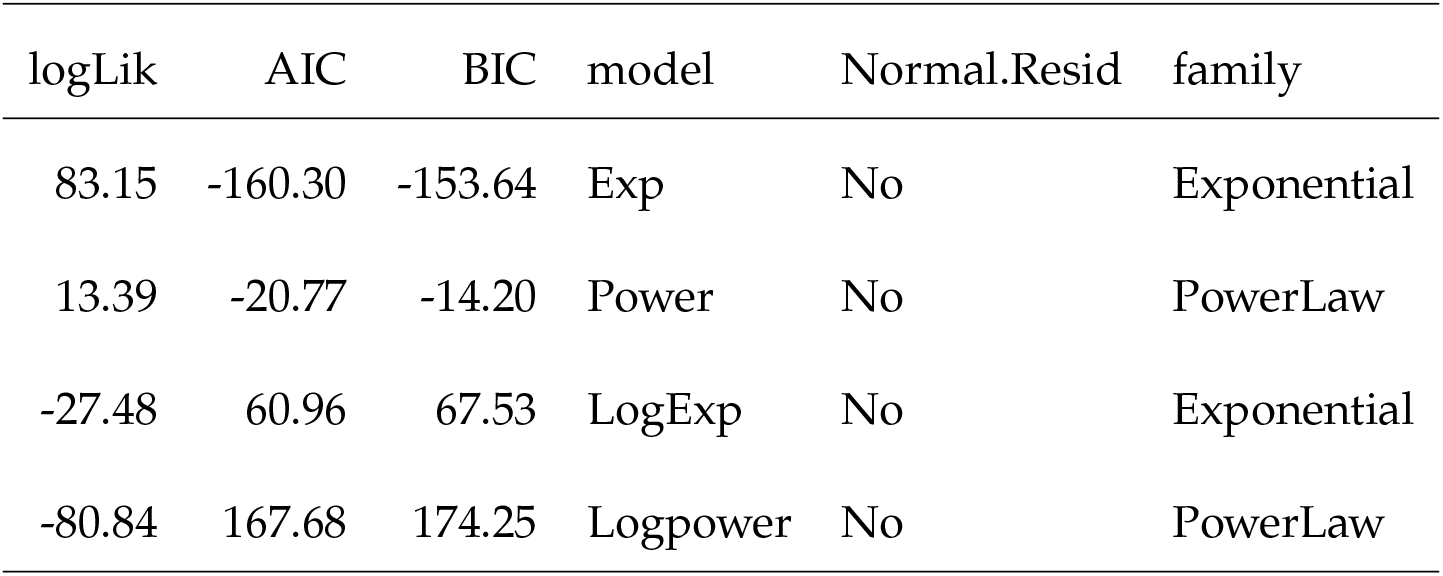
Model parameters and normal distribution tests.

| logLik | AIC | BIC | model | Normal.Resid | family |
| --- | --- | --- | --- | --- | --- |
| 83.15 | -160.30 | -153.64 | Exp | No | Exponential |
| 13.39 | -20.77 | -14.20 | Power | No | PowerLaw |
| -27.48 | 60.96 | 67.53 | LogExp | No | Exponential |
| -80.84 | 167.68 | 174.25 | Logpower | No | PowerLaw |

**Figure 5:**
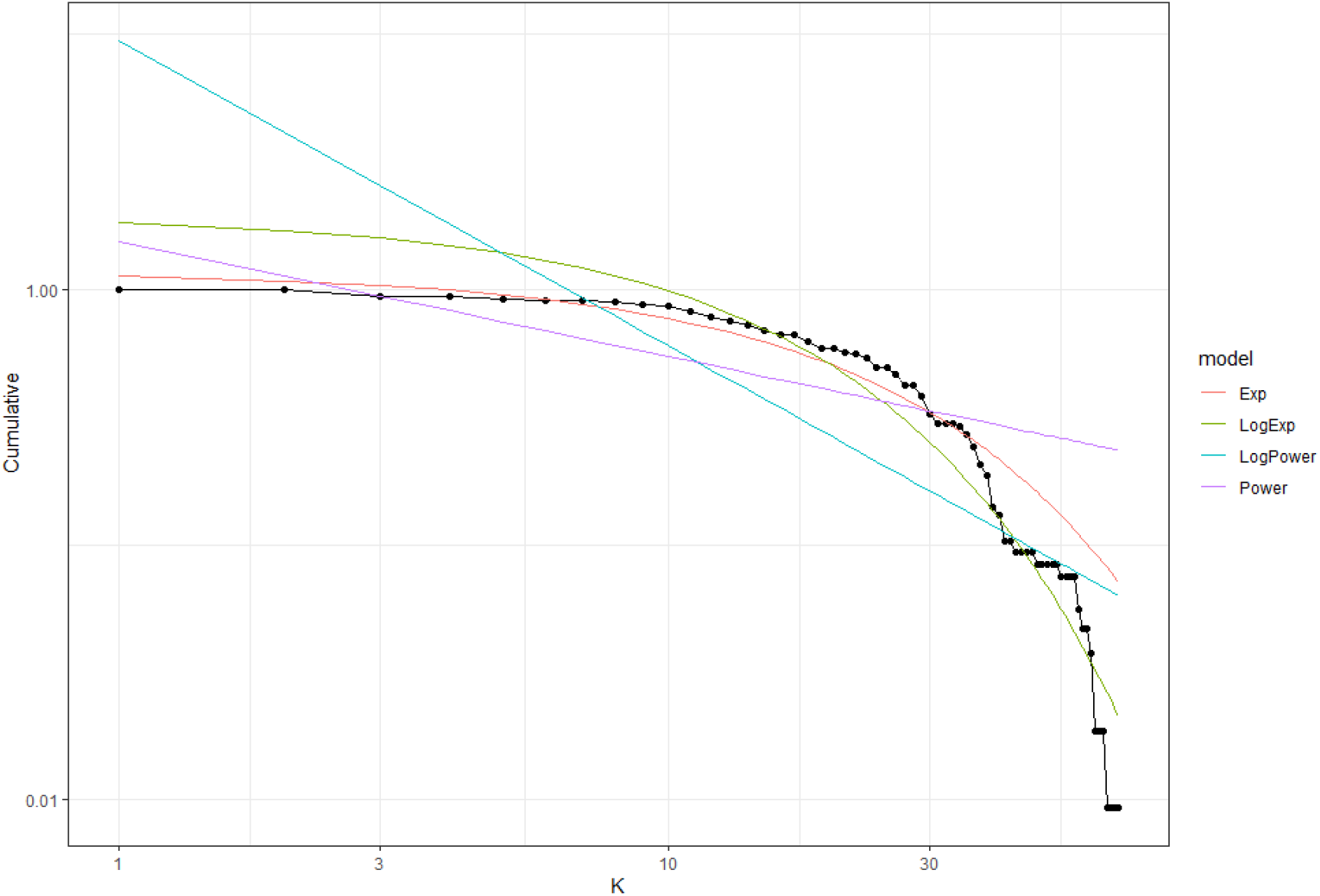
Cumulative probability distribution for a given degree (k) using the DegreeDistribution function. The plot shows two different model fits (lines). Note that since the fitted lines are regression models, their predicted values can sometimes start in values over one. Dots are the observed values

**Table 4:** Statistical parameters of the models.

| term | estimate | std.error | statistic | p.value | model |
| --- | --- | --- | --- | --- | --- |
| c | 2.25 | 0.38 | 5.84 | 0 | LogPower |
| Beta | -1.20 | 0.11 | -10.44 | 0 | LogPower |
| Beta | -0.45 | 0.04 | -11.17 | 0 | Power |
| c | 1.54 | 0.15 | 10.27 | 0 | Power |
| c | 0.67 | 0.09 | 7.25 | 0 | LogExp |
| Lambda | -0.07 | 0.00 | -28.41 | 0 | LogExp |
| Lambda | -0.04 | 0.00 | -26.98 | 0 | Exp |
| c | 0.16 | 0.03 | 6.63 | 0 | Exp |

In our example, the best model is the exponential degree distribution obtained from non-linear regressions (NLR) with an AIC = −160.30 (see Table 3). If we calculate the difference between the AIC value obtained from NLR (Exp model) with the AIC value obtained from LR (LogExp) (−160.30 - 60.96 = −221.26), it is < −2, which means that we proceed with the results obtained from NLR. Thus, the intertidal food web follows an exponential degree distribution (Figure 5).

### Extinction Thresholds - Using Weighted Networks

Biological interactions may be expressed either as present or absent, or quantified via a host of measures such as interaction frequency (González-Castro *et al.*, 2021), diet composition proportion (Cuff *et al.*, 2021), or handling time of food items (Sentis *et al.*, 2021), among others. Such weighted interactions are used to create weighted ecological networks and establish a spectrum of importance of interaction partners for each node. For example, the loss of a prey comprising 70% of a predator diet constitutes a much greater risk to its own continued existence than the loss of a prey item accounting for only 5%.

Using the argument `IS` (short for “interaction strength”) in the `SimulateExtinctions` and `RandomExtinctons` functions, users may define what proportion of original interaction strength each node is required to retain before being considered secondarily extinct. The default value is 0, denoting that a node has to become fully unconnected from the network to be considered secondarily extinct. The `IS` The argument may be used to either set a global extinction threshold or index local extinction thresholds for each individual node. Here, we demonstrate the extinction threshold argument with a global threshold of 0.5 - each node goes secondarily extinct when it looses more than 50% of its original interaction strength. To do so, we use the “`chilean_weighted`” data object supplied with the *NetworkExtinction* package (see code chunk 8 in the supplementary material). Figure 6 shows clearly how much more drastic the accumulation of secondary extinctions turns out when accounting for extinction thresholds particularly when compared to Figure 4.

**Figure 6:**
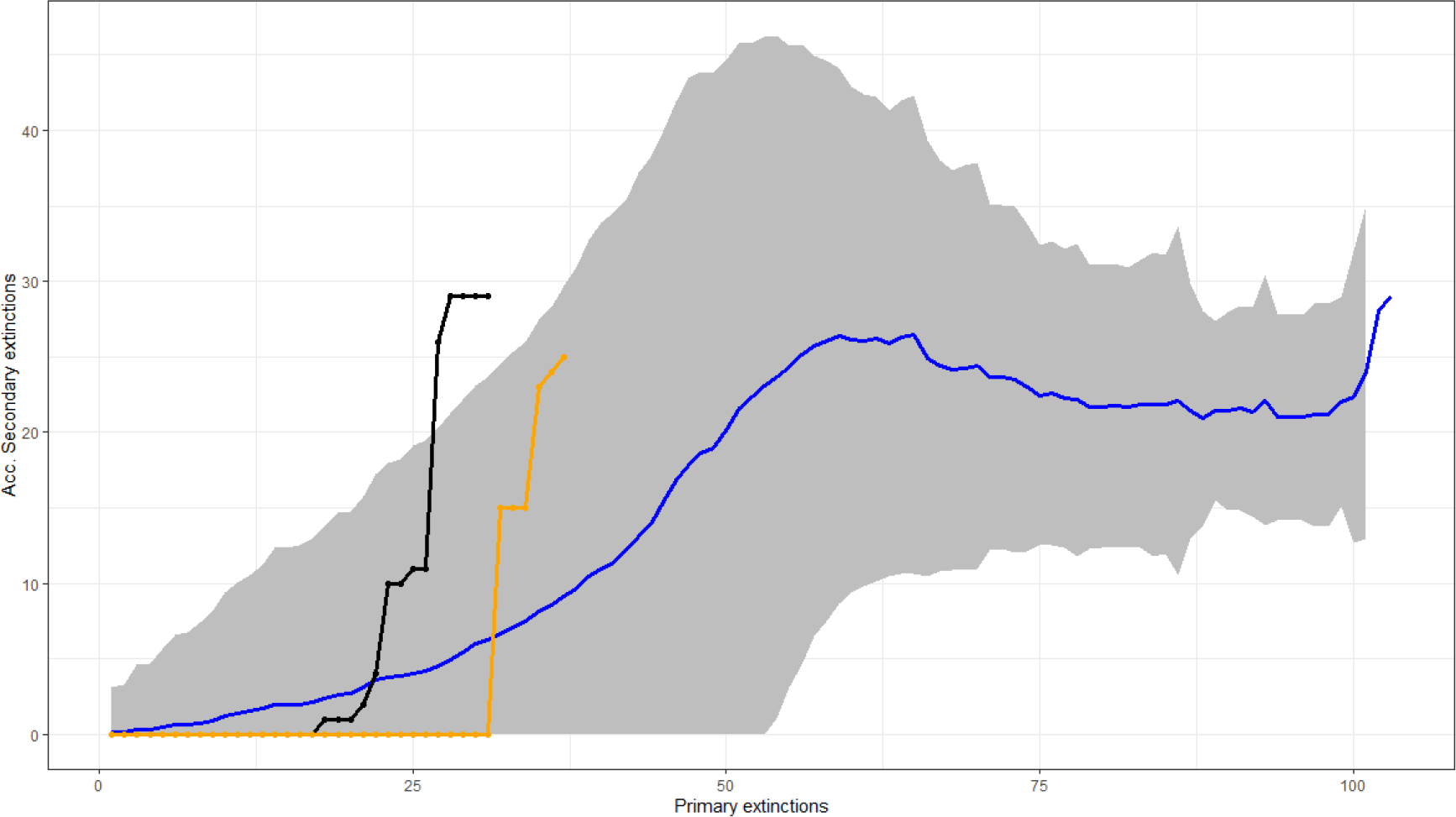
Comparison of the cumulative secondary extinctions after each removal step (Primary extinctions) between the random (Null hypothesis) and *“Mostconnected”* (Observed) deletion sequence in the weighted intertidal food web assuming an extinction threshold of 0.5. The blue line is the average (±95%CI [grey area]) of secondary extinctions of the null model and the black line following the dots represents the secondary extinctions of the observed model. The orange line represents the observed model assuming an extinction threshold of 0 (Figure 4).

To highlight the relevance of the chosen extinction threshold to the output obtained by the *NetworkExtinction* package, we have run the `SimulateExtinctions` function with the “`Mostconnected`” method for all possible values of `IS` between it’s minimum of 0 and maximum of 1 in steps of 0.01. We extracted the primary removal step at which the entire network had become unconnected/fully extinct and visualise the results in Figure 7 which shows the drastically increased rate of secondary extinctions as `IS` approaches 1.

**Figure 7:**
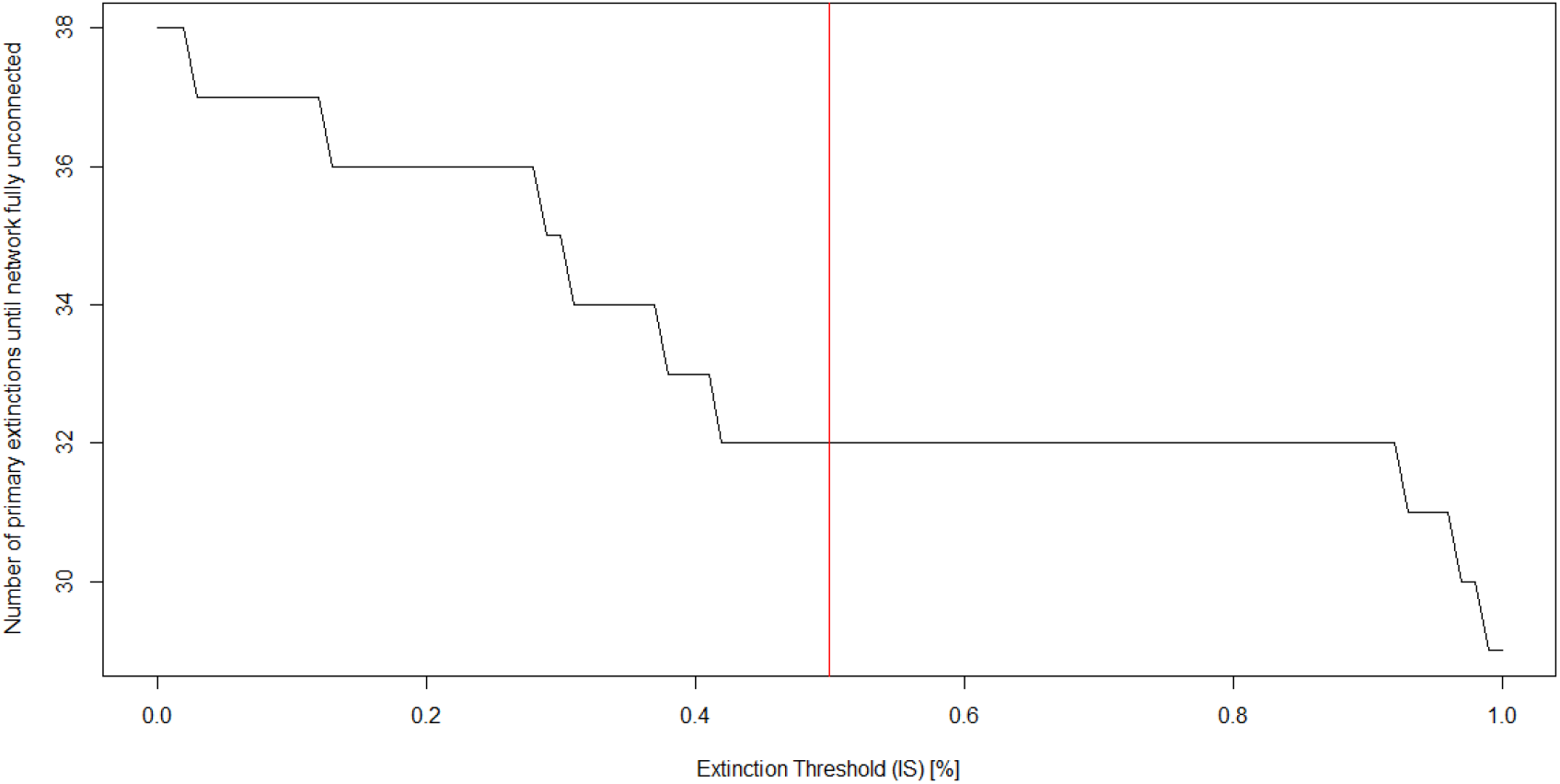
Network robustness (network of primary extinctions required to produce total disconnection of the network) over the the value space of the extinction threshold parameter. The red line indicates the extinction simulation depicted in Figure 6. See code chunk 9 in the supplementary material for this computation.

### Realising Rewiring Potential - Escape From Cascades

So far, we have demonstrated the use of the *NetworkExtinction* package under the assumption of static links. However, this assumption rarely holds in nature, where networks have been demonstrated to be capable of rewiring to new or pre-existing partners (Bartley *et al.*, 2019; Schleuning *et al.*, 2016). We have implemented functionality to account for rewiring potential in the *NetworkExtinction* package through three optional arguments to the `SimulateExtinctions` and `RandomExtinctons` functions. These are:

- `RewiringDist` - this must be a matrix of the same dimensions as the adjacency matrix defining the `Network` argument and contain either species-(dis)similarities or rewiring probabilities.
- `Rewiring` - this argument must be a function that calculates rewiring probabilities from the species-(dis)similarities stored in the `RewiringDist` object. This argument can be defined much like the `IS` argument either globally or separately for each species.
- `RewiringProb` - this global threshold determines what level of rewiring probability must be exceeded for rewiring potential to be realised.

Following a primary extinction, the *NetworkExtinction* package identifies all links which are being lost due to the removal of the primary extinction node. Then it identifies all the nodes involved in these interactions that still remain in the network. Calculating rewiring probability from `RewiringDist` matrix using the `Rewiring` function, the *NetworkExtinction* package then identifies which potential rewiring options are realised by evaluating the computed rewiring probabilities against the `RewiringProb` threshold. Any of the previously identified links for whom a realization of rewiring potential has been identified are then transferred to the new interaction partner. If there exists a pre-existing link between these two, the rewired link’s weight is addedd to the pre-existing link’s weight.

Here, we demonstrate the use of the *NetworkExtinction* package with already identified rewiring probabilities thus specifying a `Rewiring` argument which simply passes the values stored in RewiringDist along to the evaluation against the `RewiringProb` argument. To identify potential links (i.e., rewiring potential), we assigned each species into functional groups and subsequently assume that a predator preying on any item of a specific functional group may also predate each other member of the same functional group. This results in a binary matrix of potential trophic interactions in the Chilean intertidal ecosystem. This data is available via the *NetworkExtinction* package as the `chilean_potential` object. See code chunk 10 in the supplementary material for the computation. As Figure 8 indicates, accounting for rewiring potential of ecological networks leads to higher network robustness and longer runs of primary extinction simulations until full network annihilation is reached. Additionally, Figure 8 highlights that realisation of rewiring potential may lead to concentration of links on a small subset of species which incur a large number of secondary extinctions when they are removed.

**Figure 8:**
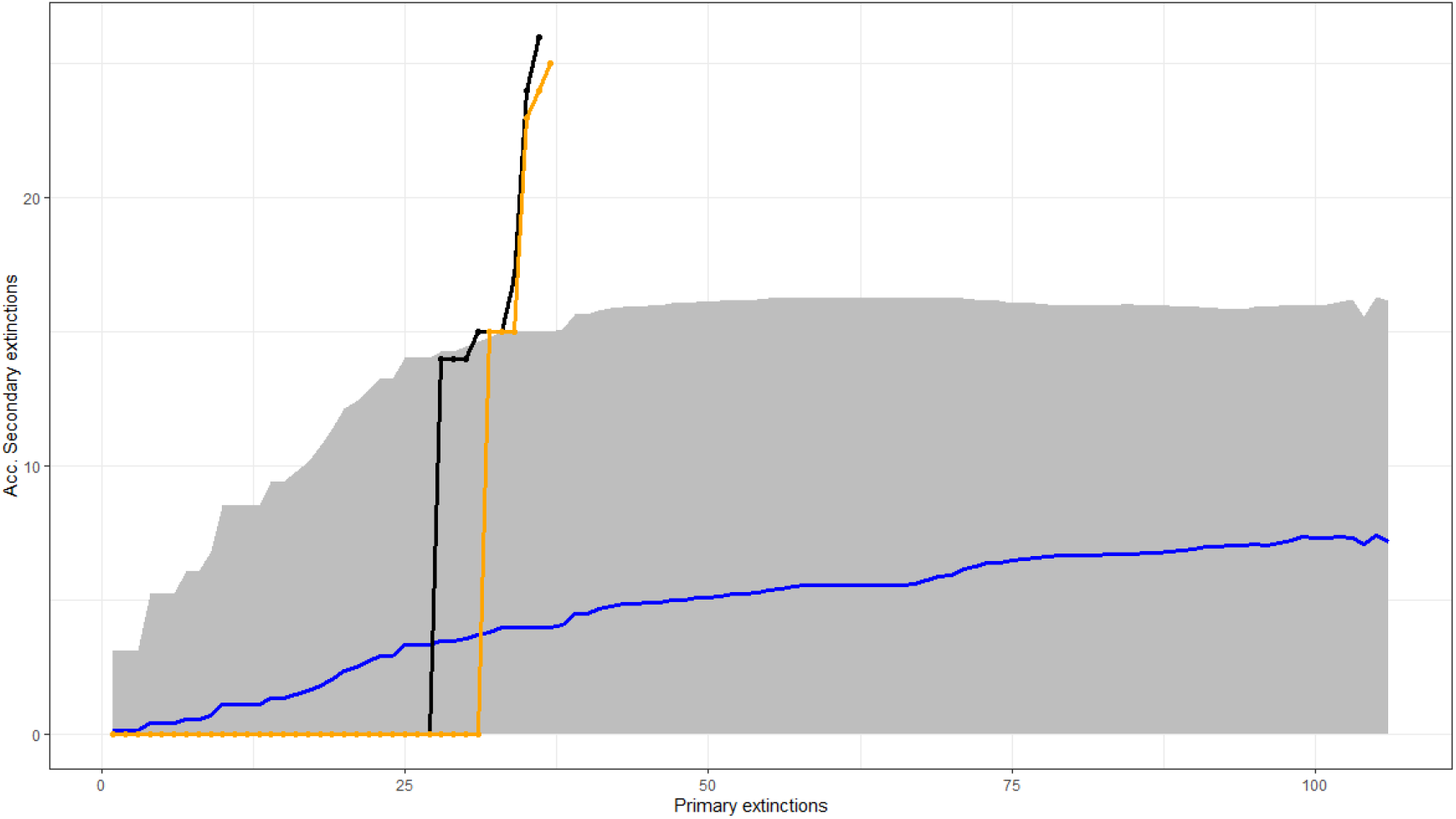
Comparison of the cumulative secondary extinctions after each removal step (Primary extinctions) between the random (Null hypothesis) and *“Mostconnected”* (Observed) deletion sequence in the intertidal food web assuming rewiring as indicated in the main text. The blue line is the average (±95%*CI* [grey area]) of secondary extinctions of the null model and the black line following the dots represents the secondary extinctions of the observed model. The orange line represents the observed model assuming no realisation of rewiring potential (Figure 4).

We realize that the implementation of the rewiring capabilities may be overly simplistic for some purposes as interactions may not be rewired wholesale, but only incrementally and split among multiple partners rather than just one rewiring partner. Nevertheless, we suggest that the capability to analyse realisation of rewiring potential in the first place represents a step-change improvement for the field of ecological network analysis and subsequent considerations of more nuanced rewiring processes may be implemented in the *NetworkExtinction* package due to its open-source nature.

## Concluding remarks

With the *NetworkExtinction* package, we have developed an easy-to-use package to visualize and assess the structure and robustness of the ecological network to different sequences of loss of species. The package lowers drastically the barrier of entry into extinction consequence forecasting models for a wide user-basis and we expect it’s applicability will be wide-ranging given the ubiquity of ecological networks.

## Acknowledgments

M.I.A. acknowledges funding from ANID-PIA/Basal FB0002, Walton Family Foundation, and to Fondecyt 3220110. P.M. acknowledges funding from Instituto de Ecología y Biodiversidad, IEB Chile (AFB-17008), FONDECYT 1200925, and ACE210010 and FB210005, BASAL funds for centers of excellence from ANID-Chile to Centro de Modelamiento Matemático (CMM). F.S.V acknowledges funding from US National Science Foundation (NSF) grants DEB-2129757 and DEB-2224915.

## Author’s contributions

M.I.A. D.C. and P.M. conceived the study. M.I.A., D.C., and E.K. wrote the code. S.A.N. provided the data for the intertidal food web. F.S.V. provided conceptual and technical support. M.I.A., D.C. and S.P.C. wrote the first draft of the manuscript. E.K. made substantial revisions to the manuscript and produced the final draft. All authors contributed to the final version of the paper.

## Data and code availability

The code for the R package can be found in the project repository (github.com/derek-corcoran-barrios/NetworkExtinction).

