## supplementary material for "NetworkExtinction: an R package to simulate extinction’s propagation and rewiring potential in ecological networks"

November 24, 2022

- **Supplementary Tables**
  - Supplementary tables : 1 - Comparison of R packages
  - Supplementary tables : 2 - Full *Mostconnected SimulateExtinctions* Results
  - Supplementary tables : 3 - Full *Ordered SimulateExtinctions* Results
  - Supplementary tables : 4 - Full *RandomExtinction* Results
- **Package Functions and Arguments**
  - Extinction Functions
  - Visualisation and Comparison Functions
  - Degree Distribution
- **Manuscript Analysis Code Chunks**

### 1 Supplementary Tables

Table 1: Comparison of the package properties between the most common package that analyze food webs with the NetworkExtinction package. We compare three categories according to the food web attributes that accept, the network metrics that calculates and the type of network analysis that performs. PP: primary producers.

| Food web attributes | Igraph | Network | Cheddar | enaR | Bipartitie | NetworkExtinction |
|---|---|---|---|---|---|---|
| Non directed networks | ✓ | ✓ | | | ✓ | |
| Directed networks | ✓ | ✓ | ✓ | ✓ | | ✓ |
| Weighted networks | ✓ | ✓ | ✓ | ✓ | ✓ | ✓ |
| Accept other node attributes (e.g. abundance, body size) | ✓ | ✓ | ✓ | ✓ | | |
| Accept environmental factors | | | ✓ | | | |
| **Food web metrics** | | | | | | |
| Calculate Number of nodes | ✓ | ✓ | ✓ | ✓ | ✓ | ✓ |
| Calculate Numbers of Links | ✓ | ✓ | ✓ | ✓ | ✓ | ✓ |
| Calculate Density of links | ✓ | ✓ | ✓ | | ✓ | ✓ |
| Calculate Connectance | | | ✓ | | ✓ | ✓ |
| Calculate other ecological index (e.g.Throphic level, omnivory level) | | | ✓ | | ✓ | |
| Calculate degree distribution | ✓ | | ✓ | | ✓ | ✓ |
| Plot degree distribution analysis | | | ✓ | | ✓ | ✓ |
| Adjust the best model of degree distribution | | | | | ✓ | ✓ |
| Use different statistical methods to adjust degree distribution | | | | | | ✓ |
| Integration with null models | | | | | ✓ | |
| **Food web analysis** | | | | | | |
| Removal of nodes | ✓ | ✓ | ✓ | | ✓ | ✓ |
| Remove a sequence of nodes with particular order | | | ✓ | | ✓ | ✓ |
| Remove nodes sequentially in order of degree | | | | | ✓ | ✓ |
| When the most connected node are removed, recalculate the new most connected node of the network | | | | | | ✓ |
| Remove nodes in a random order | | | | | ✓ | ✓ |
| Remove the secondary extincted nodes | | | | | | ✓ |
| When secondary removals occur, it is never in the PP | | | | | ✓ | ✓ |
| Quantifies the secondary extinctions of the network | | | ✓ | | ✓ | ✓ |
| Visualization of removal analysis | | | | | | ✓ |
| Statistic comparison between different removal analysis | | | | | | ✓ |
| Allow for rewiring | | | | | | ✓ |

Table 2: Full results of the *SimulateExtinctions* function with the *Mostconnected* method for the intertidal food web. Spp: node position of the extinct species, S: richness, L: number of links, C: connectance, Link_density: link density (L/S), SecExt: secondary extinctions, Pred_release: predation release, Iso_nodes: isolated nodes, AccSecExt: cumulative number of secondary extinctions, NumExt: cumulative number of primary extinctions, TotalExt: number of total extinctions.

| Spp | S | L | C | Link_density | Modularity | SecExt | Pred_release | Iso_nodes | AccSecExt | NumExt | TotalExt |
|---|---|---|---|---|---|---|---|---|---|---|---|
| 15 | 106 | 1314 | 0.12 | 12.40 | 0.01 | 0 | 0 | 0 | 0 | 1 | 1 |
| 13 | 105 | 1252 | 0.11 | 11.92 | 0.01 | 0 | 0 | 0 | 0 | 2 | 2 |
| 4 | 104 | 1192 | 0.11 | 11.46 | 0.01 | 0 | 0 | 0 | 0 | 3 | 3 |
| 12 | 103 | 1132 | 0.17 | 10.99 | 0.02 | 0 | 0 | 0 | 0 | 4 | 4 |
| 7 | 102 | 1073 | 0.10 | 10.51 | 0.02 | 0 | 0 | 0 | 0 | 5 | 5 |
| 8 | 101 | 1016 | 0.10 | 10.06 | 0.02 | 0 | 1 | 1 | 0 | 6 | 6 |
| 6 | 100 | 960 | 0.10 | 9.60 | 0.02 | 0 | 1 | 1 | 0 | 7 | 7 |
| 11 | 99 | 904 | 0.09 | 9.13 | 0.02 | 0 | 1 | 1 | 0 | 8 | 8 |
| 5 | 98 | 852 | 0.09 | 8.70 | 0.02 | 0 | 6 | 6 | 0 | 9 | 9 |
| 14 | 97 | 805 | 0.09 | 8.30 | 0.03 | 0 | 7 | 7 | 0 | 10 | 10 |
| 27 | 96 | 764 | 0.08 | 7.96 | 0.03 | 0 | 11 | 11 | 0 | 11 | 11 |
| 30 | 95 | 725 | 0.08 | 7.63 | 0.04 | 0 | 11 | 11 | 0 | 12 | 12 |
| 31 | 94 | 686 | 0.08 | 7.30 | 0.02 | 0 | 11 | 11 | 0 | 13 | 13 |
| 32 | 93 | 647 | 0.08 | 7.00 | 0.03 | 0 | 15 | 15 | 0 | 14 | 14 |
| 20 | 92 | 609 | 0.07 | 6.62 | 0.03 | 0 | 15 | 15 | 0 | 15 | 15 |
| 16 | 91 | 572 | 0.07 | 6.29 | 0.11 | 0 | 15 | 15 | 0 | 16 | 16 |
| 17 | 90 | 535 | 0.07 | 5.94 | 0.17 | 0 | 15 | 15 | 0 | 17 | 17 |
| 18 | 89 | 498 | 0.06 | 5.60 | 0.14 | 0 | 18 | 18 | 0 | 18 | 18 |
| 21 | 88 | 462 | 0.06 | 5.25 | 0.21 | 0 | 18 | 18 | 0 | 19 | 19 |
| 22 | 87 | 426 | 0.06 | 4.90 | 0.23 | 0 | 18 | 18 | 0 | 20 | 20 |
| 24 | 86 | 390 | 0.05 | 4.54 | 0.26 | 0 | 18 | 18 | 0 | 21 | 21 |
| 25 | 85 | 354 | 0.05 | 4.17 | 0.33 | 0 | 18 | 18 | 0 | 22 | 22 |
| 23 | 84 | 319 | 0.05 | 3.80 | 0.35 | 0 | 18 | 18 | 0 | 23 | 23 |
| 29 | 83 | 284 | 0.04 | 3.42 | 0.37 | 0 | 21 | 21 | 0 | 24 | 24 |
| 9 | 82 | 250 | 0.04 | 3.05 | 0.37 | 0 | 21 | 21 | 0 | 25 | 25 |

continuation of table 2.

| Spp | S | L | C | Link_density | Modularity | SecExt | Pred_release | Iso_nodes | AccSecExt | NumExt | TotalExt |
|---|---|---|---|---|---|---|---|---|---|---|---|
| 26 | 81 | 219 | 0.03 | 2.70 | 0.35 | 0 | 26 | 26 | 0 | 26 | 26 |
| 2 | 80 | 196 | 0.03 | 2.45 | 0.38 | 0 | 27 | 26 | 0 | 27 | 27 |
| 10 | 79 | 176 | 0.03 | 2.23 | 0.40 | 0 | 27 | 26 | 0 | 28 | 28 |
| 98 | 78 | 159 | 0.03 | 2.04 | 0.37 | 0 | 27 | 26 | 0 | 29 | 29 |
| 99 | 77 | 142 | 0.02 | 1.84 | 0.32 | 0 | 27 | 26 | 0 | 30 | 30 |
| 3 | 76 | 125 | 0.02 | 1.65 | 0.35 | 0 | 27 | 26 | 0 | 31 | 31 |
| 95 | 75 | 110 | 0.02 | 1.47 | 0.47 | 15 | 27 | 26 | 15 | 32 | 47 |
| 100 | 59 | 30 | 0.01 | 0.51 | 0.54 | 0 | 32 | 32 | 15 | 33 | 48 |
| 106 | 58 | 19 | 0.01 | 0.33 | 0.45 | 0 | 41 | 41 | 15 | 34 | 49 |
| 67 | 57 | 10 | 0.00 | 0.18 | 0.40 | 8 | 41 | 47 | 23 | 35 | 58 |
| 107 | 48 | 1 | 0.00 | 0.02 | 0 | 1 | 46 | 46 | 24 | 36 | 60 |
| 33 | 46 | 0 | 0.00 | 0.00 | 0 | 1 | 45 | 46 | 25 | 37 | 62 |

Table 3: Full results of the *SimulateExtinctions* function with the *Ordered* method for the intertidal food web. See column names in Table S2.

| Spp | S | L | C | Link_density | Modularity | SecExt | Pred_release | Iso_nodes | AccSecExt | NumExt | TotalExt |
|---|---|---|---|---|---|---|---|---|---|---|---|
| 67 | 106 | 1345 | 0.12 | 12.69 | 1.00E-04 | 0 | 0 | 0 | 0 | 1 | 1 |
| 37 | 105 | 1309 | 0.12 | 12.47 | 0 | 0 | 0 | 0 | 0 | 2 | 2 |
| 69 | 104 | 1275 | 0.12 | 12.26 | 0 | 0 | 0 | 0 | 0 | 3 | 3 |
| 80 | 103 | 1244 | 0.12 | 12.08 | 0 | 0 | 0 | 0 | 0 | 4 | 4 |
| 93 | 102 | 1213 | 0.12 | 11.89 | 0 | 0 | 0 | 0 | 0 | 5 | 5 |
| 38 | 101 | 1183 | 0.12 | 11.71 | 0 | 0 | 0 | 0 | 0 | 6 | 6 |
| 46 | 100 | 1153 | 0.12 | 11.53 | 0 | 0 | 0 | 0 | 0 | 7 | 7 |
| 79 | 99 | 1123 | 0.11 | 11.34 | 0 | 0 | 0 | 0 | 0 | 8 | 8 |
| 84 | 98 | 1093 | 0.11 | 11.15 | 0 | 0 | 0 | 0 | 0 | 9 | 9 |
| 94 | 97 | 1063 | 0.11 | 10.96 | 0 | 0 | 0 | 0 | 0 | 10 | 10 |
| 52 | 96 | 1034 | 0.11 | 10.77 | 0 | 0 | 0 | 0 | 0 | 11 | 11 |
| 53 | 95 | 1005 | 0.11 | 10.58 | 0 | 0 | 0 | 0 | 0 | 12 | 12 |
| 90 | 94 | 976 | 0.11 | 10.38 | 0 | 0 | 0 | 0 | 0 | 13 | 13 |
| 62 | 93 | 947 | 0.11 | 10.18 | 0 | 0 | 0 | 0 | 0 | 14 | 14 |
| 40 | 92 | 920 | 0.11 | 10 | 0 | 0 | 0 | 0 | 0 | 15 | 15 |
| 42 | 91 | 893 | 0.11 | 9.81 | 0 | 0 | 0 | 0 | 0 | 16 | 16 |
| 43 | 90 | 866 | 0.11 | 9.62 | 0 | 0 | 0 | 0 | 0 | 17 | 17 |
| 44 | 89 | 839 | 0.11 | 9.43 | 0 | 1 | 0 | 1 | 1 | 18 | 19 |
| 60 | 87 | 812 | 0.11 | 9.33 | 0 | 0 | 0 | 0 | 1 | 19 | 20 |
| 59 | 86 | 786 | 0.11 | 9.14 | 0 | 0 | 0 | 0 | 1 | 20 | 21 |
| 85 | 85 | 760 | 0.11 | 8.94 | 0 | 0 | 0 | 0 | 1 | 21 | 22 |
| 87 | 84 | 734 | 0.1 | 8.74 | 0 | 0 | 0 | 0 | 1 | 22 | 23 |
| 55 | 83 | 712 | 0.1 | 8.58 | 0 | 0 | 0 | 0 | 1 | 23 | 24 |
| 72 | 82 | 689 | 0.1 | 8.4 | 0 | 0 | 0 | 0 | 1 | 24 | 25 |
| 91 | 81 | 666 | 0.1 | 8.22 | 0 | 0 | 0 | 0 | 1 | 25 | 26 |
| 74 | 80 | 645 | 0.1 | 8.06 | 0 | 0 | 0 | 0 | 1 | 26 | 27 |
| 68 | 79 | 624 | 0.1 | 7.9 | 0 | 0 | 0 | 0 | 1 | 27 | 28 |
| 75 | 78 | 605 | 0.1 | 7.76 | 0 | 6 | 0 | 0 | 7 | 28 | 35 |
| 65 | 71 | 521 | 0.1 | 7.34 | 0.04 | 0 | 0 | 0 | 7 | 29 | 36 |
| 88 | 70 | 503 | 0.1 | 7.19 | 0.04 | 0 | 0 | 0 | 7 | 30 | 37 |
| 33 | 69 | 487 | 0.1 | 7.06 | 0.04 | 0 | 0 | 0 | 7 | 31 | 38 |

continuation of table 3.

| Spp | S | L | C | Link_density | Modularity | SecExt | Pred_release | Iso_nodes | AccSecExt | NumExt | TotalExt |
|---|---|---|---|---|---|---|---|---|---|---|---|
| 34 | 68 | 471 | 0.1 | 6.93 | 0.04 | 0 | 0 | 0 | 7 | 32 | 39 |
| 45 | 67 | 455 | 0.1 | 6.79 | 0.04 | 0 | 0 | 0 | 7 | 33 | 40 |
| 20 | 67 | 455 | 0.1 | 6.79 | 0.04 | 0 | 0 | 0 | 7 | 34 | 41 |
| 29 | 66 | 436 | 0.1 | 6.61 | 0.04 | 0 | 0 | 0 | 7 | 35 | 42 |
| 64 | 65 | 422 | 0.1 | 6.49 | 0.04 | 1 | 0 | 0 | 8 | 36 | 44 |
| 66 | 63 | 400 | 0.1 | 6.35 | 0.05 | 0 | 0 | 0 | 8 | 37 | 45 |
| 95 | 62 | 391 | 0.1 | 6.31 | 0 | 9 | 0 | 0 | 17 | 38 | 55 |
| 51 | 52 | 337 | 0.12 | 6.48 | 0 | 3 | 0 | 0 | 20 | 39 | 59 |
| 81 | 48 | 307 | 0.13 | 6.4 | 0 | 0 | 0 | 0 | 20 | 40 | 60 |
| 36 | 48 | 307 | 0.13 | 6.4 | 0 | 0 | 0 | 0 | 20 | 41 | 61 |
| 47 | 47 | 294 | 0.13 | 6.26 | 0 | 0 | 0 | 0 | 20 | 42 | 62 |
| 48 | 46 | 281 | 0.13 | 6.11 | 0 | 0 | 0 | 0 | 20 | 43 | 63 |
| 49 | 45 | 268 | 0.13 | 5.96 | 0 | 3 | 0 | 0 | 23 | 44 | 67 |
| 15 | 41 | 237 | 0.14 | 5.78 | 0 | 0 | 0 | 0 | 23 | 45 | 68 |
| 21 | 41 | 237 | 0.14 | 5.78 | 0 | 0 | 0 | 0 | 23 | 46 | 69 |
| 24 | 41 | 237 | 0.14 | 5.78 | 0 | 0 | 0 | 0 | 23 | 47 | 70 |
| 54 | 40 | 227 | 0.14 | 5.68 | 0 | 1 | 0 | 0 | 24 | 48 | 72 |
| 83 | 38 | 205 | 0.14 | 5.39 | 0 | 3 | 0 | 0 | 27 | 49 | 76 |
| 7 | 34 | 173 | 0.15 | 5.09 | 0 | 1 | 0 | 1 | 28 | 50 | 78 |
| 9 | 33 | 173 | 0.16 | 5.24 | 0 | 0 | 0 | 0 | 28 | 51 | 79 |
| 12 | 32 | 154 | 0.15 | 4.81 | 0 | 0 | 0 | 0 | 28 | 52 | 80 |
| 13 | 31 | 134 | 0.14 | 4.32 | 0 | 0 | 0 | 0 | 28 | 53 | 81 |
| 22 | 31 | 134 | 0.14 | 4.32 | 0 | 0 | 0 | 0 | 28 | 54 | 82 |
| 25 | 31 | 134 | 0.14 | 4.32 | 0 | 0 | 0 | 0 | 28 | 55 | 83 |
| 39 | 31 | 134 | 0.14 | 4.32 | 0 | 0 | 0 | 0 | 28 | 56 | 84 |
| 70 | 30 | 127 | 0.14 | 4.23 | 0 | 0 | 0 | 0 | 28 | 57 | 85 |
| 73 | 29 | 120 | 0.14 | 4.14 | 0 | 0 | 0 | 0 | 28 | 58 | 86 |
| 89 | 28 | 113 | 0.14 | 4.04 | 0 | 0 | 0 | 0 | 28 | 59 | 87 |
| 6 | 27 | 97 | 0.13 | 3.59 | 0 | 0 | 0 | 0 | 28 | 60 | 88 |

Table 4: Summarised results of the *RandomExtinction* function for the intertidal food web, showing the first and last three rows. NumExt: cumulative number of primary extinctions, AccSecExt_95CI: cumulative 95% of confidence intervals of the secondary extinctions among all the simulations performed, AccSecExt_mean: cumulative average of the secondary extinctions among all the simulations performed, Upper & Lower: lower and upper limit of the [mean + 95% CI], respectively.

| NumExt | AccSecExt_95CI | AccSecExt_mean | Upper | Lower |
|---:|---:|---:|---:|---:|
| 1 | 0 | 0 | 0 | 0 |
| 2 | 0 | 0 | 0 | 0 |
| 3 | 4.14 | 0.3 | 4.44 | 0 |
| 4 | 5.7 | 0.59 | 6.29 | 0 |
| 5 | 5.7 | 0.59 | 6.29 | 0 |
| 6 | 5.7 | 0.59 | 6.29 | 0 |
| 7 | 6.36 | 0.74 | 7.1 | 0 |
| 8 | 6.94 | 0.89 | 7.83 | 0 |
| 9 | 6.94 | 0.89 | 7.83 | 0 |
| 10 | 6.94 | 0.89 | 7.83 | 0 |
| 11 | 6.94 | 0.89 | 7.83 | 0 |
| 12 | 7.27 | 1.01 | 8.28 | 0 |
| 13 | 7.77 | 1.16 | 8.93 | 0 |
| 14 | 8.1 | 1.29 | 9.39 | 0 |
| 15 | 8.1 | 1.29 | 9.39 | 0 |
| 16 | 8.37 | 1.41 | 9.78 | 0 |
| 17 | 8.9 | 1.66 | 10.56 | 0 |
| 18 | 8.9 | 1.66 | 10.56 | 0 |
| 19 | 8.9 | 1.66 | 10.56 | 0 |
| 20 | 9.27 | 1.81 | 11.08 | 0 |
| 21 | 9.47 | 1.93 | 11.4 | 0 |
| 22 | 9.47 | 1.93 | 11.4 | 0 |
| 23 | 9.67 | 2.05 | 11.72 | 0 |
| 24 | 10.11 | 2.31 | 12.42 | 0 |
| 25 | 10.51 | 2.57 | 13.08 | 0 |
| 26 | 10.73 | 2.71 | 13.44 | 0 |
| 27 | 10.73 | 2.71 | 13.44 | 0 |
| 28 | 11.09 | 3.13 | 14.22 | 0 |

continuation of table 4.

| NumExt | AccSecExt_95CI | AccSecExt_mean | Upper | Lower |
|---:|---:|---:|---:|---:|
| 29 | 11.19 | 3.16 | 14.35 | 0 |
| 30 | 11.26 | 3.26 | 14.52 | 0 |
| 31 | 11.34 | 3.37 | 14.71 | 0 |
| 32 | 11.34 | 3.37 | 14.71 | 0 |
| 33 | 11.52 | 3.6 | 15.12 | 0 |
| 34 | 11.53 | 3.82 | 15.35 | 0 |
| 35 | 11.55 | 3.84 | 15.39 | 0 |
| 36 | 11.55 | 3.84 | 15.39 | 0 |
| 37 | 11.57 | 3.93 | 15.5 | 0 |
| 38 | 11.69 | 4.07 | 15.76 | 0 |
| 39 | 11.79 | 4.28 | 16.07 | 0 |
| 40 | 11.82 | 4.29 | 16.11 | 0 |
| 41 | 11.88 | 4.59 | 16.47 | 0 |
| 42 | 11.89 | 4.6 | 16.49 | 0 |
| 43 | 11.89 | 4.6 | 16.49 | 0 |
| 44 | 11.88 | 4.77 | 16.65 | 0 |
| 45 | 11.86 | 4.85 | 16.71 | 0 |
| 46 | 11.87 | 4.86 | 16.73 | 0 |
| 47 | 11.91 | 4.98 | 16.89 | 0 |
| 48 | 11.91 | 5.08 | 16.99 | 0 |
| 49 | 11.92 | 5.19 | 17.11 | 0 |
| 50 | 11.94 | 5.2 | 17.14 | 0 |
| 51 | 11.9 | 5.27 | 17.17 | 0 |
| 52 | 11.85 | 5.44 | 17.29 | 0 |
| 53 | 11.82 | 5.53 | 17.35 | 0 |
| 54 | 11.7 | 5.77 | 17.47 | 0 |
| 55 | 11.61 | 5.93 | 17.54 | 0 |
| 56 | 11.58 | 6.03 | 17.61 | 0 |
| 57 | 11.52 | 6.1 | 17.62 | 0 |
| 58 | 11.52 | 6.1 | 17.62 | 0 |
| 59 | 11.55 | 6.11 | 17.66 | 0 |
| 60 | 11.48 | 6.21 | 17.69 | 0 |
| 61 | 11.29 | 6.44 | 17.73 | 0 |
| 62 | 11.19 | 6.51 | 17.7 | 0 |
| 63 | 11.04 | 6.63 | 17.67 | 0 |

continuation of table 4.

| NumExt | AccSecExt_95CI | AccSecExt_mean | Upper | Lower |
|---:|---:|---:|---:|---:|
| 64 | 11.04 | 6.64 | 17.68 | 0 |
| 65 | 10.97 | 6.72 | 17.69 | 0 |
| 66 | 10.95 | 6.73 | 17.68 | 0 |
| 67 | 10.85 | 6.79 | 17.64 | 0 |
| 68 | 10.74 | 6.87 | 17.61 | 0 |
| 69 | 10.66 | 6.91 | 17.57 | 0 |
| 70 | 10.66 | 6.91 | 17.57 | 0 |
| 71 | 10.66 | 6.91 | 17.57 | 0 |
| 72 | 10.58 | 6.96 | 17.54 | 0 |
| 73 | 10.5 | 7.07 | 17.57 | 0 |
| 74 | 10.5 | 7.07 | 17.57 | 0 |
| 75 | 10.42 | 7.11 | 17.53 | 0 |
| 76 | 10.35 | 7.17 | 17.52 | 0 |
| 77 | 10.35 | 7.17 | 17.52 | 0 |
| 78 | 10.26 | 7.22 | 17.48 | 0 |
| 79 | 10.26 | 7.22 | 17.48 | 0 |
| 80 | 10.18 | 7.35 | 17.53 | 0 |
| 81 | 10.09 | 7.41 | 17.5 | 0 |
| 82 | 10.09 | 7.41 | 17.5 | 0 |
| 83 | 10.09 | 7.41 | 17.5 | 0 |
| 84 | 10.06 | 7.42 | 17.48 | 0 |
| 85 | 10 | 7.45 | 17.45 | 0 |
| 86 | 9.88 | 7.5 | 17.38 | 0 |
| 87 | 9.86 | 7.52 | 17.38 | 0 |
| 88 | 9.86 | 7.55 | 17.41 | 0 |
| 89 | 9.84 | 7.57 | 17.41 | 0 |
| 90 | 9.89 | 7.65 | 17.54 | 0 |
| 91 | 9.89 | 7.65 | 17.54 | 0 |
| 92 | 9.81 | 7.74 | 17.55 | 0 |
| 93 | 9.69 | 7.83 | 17.51 | 0 |
| 94 | 9.68 | 7.89 | 17.57 | 0 |
| 95 | 9.63 | 7.97 | 17.6 | 0 |
| 96 | 9.53 | 8.06 | 17.59 | 0 |
| 97 | 9.54 | 8.14 | 17.68 | 0 |
| 98 | 9.53 | 8.15 | 17.68 | 0 |

continuation of table 4.

| NumExt | AccSecExt_95CI | AccSecExt_mean | Upper | Lower |
|---:|---:|---:|---:|---:|
| 99 | 9.57 | 8.24 | 17.81 | 0 |
| 100 | 9.47 | 8.35 | 17.82 | 0 |
| 101 | 9.52 | 8.51 | 18.03 | 0 |
| 102 | 9.1 | 8.86 | 17.96 | 0 |
| 103 | 9.45 | 9.23 | 18.67 | 0 |
| 104 | 9.59 | 9.32 | 18.9 | 0 |
| 105 | 10.15 | 9.46 | 19.61 | 0 |
| 106 | 9.59 | 10.18 | 19.78 | 0.59 |

### Package Functions and Arguments

#### *Extinction Functions*

##### *SimulateExtinctions*

This function remove nodes using either of the following two deletion sequences options: 1) based on species' degree (*Mostconnected method*), 2) a user-defined order (*Ordered method*).

The *Mostconnected method* sorts the nodes (species) from the most to the least connected one, considering their total degree (in-degrees plus out-degrees), and storing the information in a vector. Then, the function calculates the following topological indices of the ecological network: species richness, number of links, network connectance, link density and modularity. Subsequently, it removes the most connected node in the network. At this point in the process of the function, all links lost from the network following this primary removal are subject to rewiring processes. These are defined through the *Rewiring*, *RewiringDist*, and *RewiringProb* arguments. *Rewiring* defines a function which translates similarity scores in the *RewiringDist* distance matrix into probabilities of rewiring. If, for any of the lost links, any rewiring probability calculated from these arguments exceeds the *RewiringProb* threshold, the links is reassigned (i.e. rewired) the novel partner with the highest rewiring probability. Note that *RewiringDist* may also represent rewiring probabilities themselves. See Kusch & Ordonez, in prep. for a demonstration of this capability. By default, rewiring processes are not considered. Next, the package recalculates the topological indexes of the ecological network and counts secondary extinctions. Extinction thresholds which govern secondary extinctions can be set either globally for the entire networks or individually for each node using the *IS* argument. The *IS* argument (percentage, expressed as a number between 0 and 1) defines how much of the original interaction strength each node may lose before being considered secondarily extinct. By default, this argument is set to 0 thus only considering a species as secondarily extinct when it's node loses all links. Next, the package removes all extinct species (primary and secondary) from the ecological network. After that removal, the algorithm recalculates the degree ranking, iterating the process until the number of links in the network is zero (Dunne & Williams, 2009; Dunne *et al.*, 2002; Sole & Montoya, 2001). In case of a tie, i.e. two node have the same degree value, the removed node is randomly chosen among them (Curtsdotter *et al.*, 2011; Valdovinos *et al.*, 2009).

In the *Ordered method* the user presets the order required to remove the nodes. After each node is removed, it calculates the secondary extinctions and the same topological indexes are calculated by the *Mostconnected* method. In this approach, the supplied order of primary extinctions is not altered as species are being removed.

This function requires the following arguments:

- *Network*: The name of the network to analyze in matrix or network class.
- *Method*: a character with the options Mostconnected or Ordered.
- *Order*: this should be NULL, unless using the Ordered method, in that case it should be a numeric vector with the node position of the species sorted in the removal order of interest. If an Order is specified, the Method is automatically considered to be Ordered.
- *NetworkType*: a character either specifying Trophic or Mutualistic network type.
- *clust.method*: a character with the clustering option to calculate modularity. The options are: cluster_infomap (default), cluster_edge_betweenness, cluster_spinglass or cluster_label_prop (Clauset *et al.*, 2004). The cluster_infomap the lightest clustering method. If you use other

options you should have a powerful computer or your RAM might not be enough, depending on network size.

- *IS*: either numeric or a named vector of numerics. Identifies the threshold of relative interaction strength which species require to not be considered secondarily extinct.
- *Rewiring*: either a function or a named vector of functions. Signifies how rewiring probabilities are calculated from the RewiringDist argument. If FALSE, no rewiring is carried out.
- *RewiringDist*: a numeric matrix of NxN dimension (N... number of nodes in Network). Contains, for example, phylogenetic or functional trait distances between nodes in Network which are used by the Rewiring argument to calculate rewiring probabilities. If *Rewiring* == *function(xx RewiringDist* is treated as a matrix storing probability of species pairs being linked to each other.
- *RewiringProb*: a numeric which identifies the threshold at which to assume rewiring potential is met.
- *verbose*: Logical. Whether to report on function progress or not.

##### *RandomExtinctions*

The `RandomExtinctions` function generates *nsim* independent random extinction sequences by bootstrapping. Each trajectory in *nsim* runs until either all species are extinct or a number of primary extinction equal to the argument *SimNum* has been simulated. After all the trajectories were simulated, the function calculates the average and standard deviation of the number of secondary extinctions through *nsim* for each removal step.

To run the `RandomExtinctions` function, the following arguments are necessary:

- *Network*: The name of the network to analyze in matrix or network class.
- *nsim*: is number of random extinction sequences chosen (10 by default).
- *parallel*: Optional argument to run the simulation in parallel trough different cores to decrease the running time. if is TRUE, it will use parallel processing, if FALSE (default) it will run sequentially.
- *ncores*: number of cores to use if using parallel processing.
- *Record*: Optional logical argument. If TRUE, records every simulation result.
- *plot*: Optional logical argument. If TRUE (default), the function returns a plot with the cumulative mean of secondary extinctions ($\pm$ 95% CI) for each removal step averaged through all the simulations.
- *SimNum*: numeric, how many nodes to register for primary extinction. By default sets all of them.
- *NetworkType*: same as in *SimulateExtinctions* function.
- *clust.method*: same as in *SimulateExtinctions* function.
- *IS*: same as in *SimulateExtinctions* function.
- *Rewiring*: same as in *SimulateExtinctions* function.
- *RewiringDist*: same as in *SimulateExtinctions* function.
- *RewiringProb*: same as in *SimulateExtinctions* function.

- *verbose*: same as in *SimulateExtinctions* function.

To make these simulations more efficient, we have implemented parallel computation for multiple random deletion sequences at the same time. Additionally, the optional argument `SimNum` controls how many nodes to include in each random deletion sequence. By default, all nodes of the network are included in each random deletion sequence. For use with user-defined deletion sequences, one may usually wish to set the `SimNum` argument to the same as the number of nodes indexed via the user-defined sequence.

#### *Visualisation and Comparison Functions*

##### *ExtinctionPlot*

The *ExtinctionPlot* function takes an output data frame from any of the extinction functions and plots the indicator of interest after each primary extinction step. By default, this function plots the number of accumulated secondary extinctions after every primary extinction, but any of the indices (e.g., link density, or predation release) can be plotted with the function by changing the *Variable* argument.

To run the `ExtinctionPlot` function, the following arguments are necessary:

- *History*: A data frame. The object returned by the *SimulateExtinction* function.

- *Variable*: is the name of the dependent variable of interest to be plotted against the number of extinctions.

*CompareExtinction*

The `CompareExtinctions` function compares the accumulative observed extinction generated by the *SimulateExtinctions* function against a null model of the expected accumulative secondary extinctions developed by the *RandomExtinctons* function.

To run the `CompareExtinctions` function, the following arguments are necessary:

- *Nullmodel*: A data frame. The object returned by the *RandomExtinctions* function.
- *Hypothesis*:A data frame. The object returned by the *SimulateExtinctions* function.

#### *DegreeDistribution*

The `DegreeDistribution` function computes the degree distribution of a network, calculating $P(k)$, i.e, the probability that a node in the network is connected to $k$ other nodes (Estrada, 2007). Then, the observed distribution is fitted to the exponential and power-law models using two different approaches, non-linear least squares (Godfrey *et al.*, 1998), and linear models on log-transformed data (Xiao *et al.*, 2011). A selected set of model fitting techniques are already available (Clauset *et al.*, 2009).

We used a two-step model selection approach to decide if the observed degree distribution of the network follows an exponential or a power-law distribution. Firstly, the function selects the best method to fitting the models (non-linear, NLR vs log-transformed linear method, LR) according to their ability to provide good estimates of precision for hypothesis testing. Secondly, after one method has been selected, the function selects the distribution (exponential vs power-law distribution) to which the data best fit. In the first step, we fit each the model of each distribution using both methods (LR and NLR) (Table **??**). Following Xiao *et al.* (2011), we used maximum likelihood fitting for testing the normal distribution of residuals, and calculated *AICc* values for each method. To do that, we compared the ΔAICc between both methods (LR and NLR) to each distribution model. If the ΔAICc between LR and LNR is larger than 2, the LR is selected. Otherwise, a model average is used between both methods using Akaike weights following (Burnham & Anderson, 2002).

After the linearized or non-linear approach has been selected, a comparison is made between the exponential vs the power-law distributions using AIC (Burnham & Anderson, 2002). By doing this, we only select the best model describing the degree distribution, since there is no way of doing model averaging between both models (for more details about the model selection see Xiao *et al.* (2011)).

To run the `DegreeDistribution` function, the following arguments are necessary:

- *Network*: The name of the network to analyze in matrix or network class.
- *scale*: a character stating if the graph is on a log-log scale ("LogLog") or arithmetic scale ("arithmetic").

### Manuscript Analysis Code Chunks

Code chunk 1: Package installation and loading.

```
R> install.packages("NetworkExtinction")

R> library(NetworkExtinction)
```

Code chunk 2: Mostconnected example with unweighted food web, with binary extinction threshold and without rewiring.

```
R> data("chilean_intertidal")

R> Mostconnected <- SimulateExtinctions(Network = chilean_intertidal,

                                        Method = "Mostconnected",

                                        NetworkType = "Trophic", # default argument

                                        IS = 0, # default argument

                                        Rewiring = FALSE # default argument

                                        )

R> Mostconnected$sims
```

Code chunk 3: Ordered example with unweighted food web, with binary extinction threshold and without rewiring.

```
R> data("chilean_intertidal")

R> Order <- order(sna::degree(chilean_intertidal, cmode = "outdegree"),

                  decreasing = TRUE)[1:60])

R> Ordered <- SimulateExtinctions(Network = chilean_intertidal,
```

```
                                Method = Ordered, Order = Order,
                                NetworkType = "Trophic", # default argument
                                IS = 0, # default argument
                                Rewiring = FALSE # default argument
                                )
R> Ordered$sims
```

Code chunk 4: Comparison of primary-only and *NetworkExtinction* simulation outcomes.

```
R> data("chilean_intertidal")
R> circlelayout <- network::network.layout.circle(chilean_intertidal)
## Primary-Only Output
R> Order_primonly <- chilean_intertidal
R> Order_primonly <- network::delete.vertices(Order_primonly, Order)
R> plot(Order_primonly, mode = "circle",
     coord = circlelayout[as.numeric(
     network::get.vertex.attribute(Order_primonly, "vertex.names")), ])
## NetworkExtinction Output
R> plot(Ordered$Network, mode = "circle",
     coord = circlelayout[as.numeric(
     network::get.vertex.attribute(Ordered$Network, "vertex.names")), ])
```

Code chunk 5: Random extinction simulations for comparison with outputs of chunk 2 and 3 (unweighted networks, binary extinction threshold, no rewiring).

```
R> data("chilean_intertidal")
R> Random_MC <- RandomExtinctions(Network = chilean_intertidal, nsim = 100,
                               parallel = TRUE, ncores = 4,
                               Record = FALSE, plot = TRUE)
R> Random_OR <- RandomExtinctions(Network = chilean_intertidal, nsim = 100,
                               SimNum = length(Order),
                               parallel = TRUE, ncores = 4,
                               Record = FALSE, plot = TRUE)
```

Code chunk 6: Visualisations of *NetworkExtinction* outputs.

```
R> ExtinctionPlot(Mostconnected$sims)
R> ExtinctionPlot(History = Mostconnected$sims, Variable = "Link_density")
R> CompareExtinctions(Nullmodel = Random_MC, Hypothesis = Mostconnected)
R> CompareExtinctions(Nullmodel = Random_OR, Hypothesis = Ordered)
```

Code chunk 7: Degree distribution function call.

```
R> DegreeDistribution(chilean_intertidal, scale = "LogLog")
```

Code chunk 8: Threshold example using a weighted network and an extinction threshold of 0.5 without rewiring.

```
R> library(ggplot2)
R> data(chilean_weighted)
## simulations
```

```
R> Mostconnected_Thresh <- SimulateExtinctions(Network = chilean_intertidal,
                                              Method = "Mostconnected",
                                              IS = 0.5)
R> Random_Thresh <- RandomExtinctions(Network = chilean_intertidal, nsim = 100,
                                     IS = 0.5, parallel = TRUE, ncores = 4)
## generating manuscript plot
R> Nullmodel <- Random_Thresh
R> Hypothesis <- Mostconnected_Thresh
R> Addmodel <- Mostconnected$sims
R> g <- ggplot(Nullmodel$sims, aes(x = NumExt, y = AccSecExt_mean)) +
  geom_ribbon(aes_string(ymin = "Lower", ymax = "Upper"), fill = "grey") +
  geom_line(aes(color = "blue"), color = "blue", size = 1.2) +
  ylab("Acc. Secondary extinctions") + xlab("Primary extinctions") + theme_bw()
R> g <- g + geom_point(data = Hypothesis$sims,
            aes(y = AccSecExt), color = "black") +
  geom_line(data = Hypothesis$sims,
            aes(y = AccSecExt, color = "black"), color = "black", size = 1.2)
R> g <- g + geom_point(data = Addmodel, aes(y = AccSecExt), color = "orange") +
  geom_line(data = Addmodel,
            aes(y = AccSecExt, color = "orange"), color = "orange", size = 1.2)
R> g
```

Code chunk 9: Identification of removal steps until full-network annihilation given varying levels of extinction thresholds for an unweighted network without rewiring.

```
R>

ThreshSeq <- seq(0, 1, 0.01)

SimSteps <- c()

for(i in ThreshSeq)  Thresh <- SimulateExtinctions(Network = chilean_intertidal,

plot(ThreshSeq, SimSteps, type = "l",

            xlab = "Extinction Threshold (IS) [%]",

            ylab = "Number of primary extinctions until network fully unconnected")

abline(v = 0.5, col = "red")
```

Code chunk 10: Rewiring example using an unweighted network with a binary extinction threshold and assuming realisation of rewiring potential to potential partners with a link probability exceeding 0.5

```
R> library(ggplot2)

R> data(chilean_intertidal)

R> data(chilean_potential)

## simulations

R> Mostconnected_Rewiring <- SimulateExtinctions(Network = chilean_intertidal,

                                            Method = "Mostconnected",

                                            Rewiring = function(x)x,

                                            RewiringDist = chilean_potential,

                                            RewiringProb = 0.5)

R> Random_Rewiring <- RandomExtinctions(Network = chilean_intertidal, nsim = 100,
```

```
                                   Method = "Mostconnected",
                                   Rewiring = function(x)x,
                                   RewiringDist = chilean_potential,
                                   RewiringProb = 0.5,
                                   parallel = TRUE, ncores = 4)
## generating manuscript plot
R> Nullmodel <- Random_Rewiring
R> Hypothesis <- Mostconnected_Rewiring
R> Addmodel <- Mostconnected$sims
R> g <- ggplot(Nullmodel$sims, aes(x = NumExt, y = AccSecExt_mean)) +
  geom_ribbon(aes_string(ymin = "Lower", ymax = "Upper"), fill = "grey") +
  geom_line(aes(color = "blue"), color = "blue", size = 1.2) +
  ylab("Acc. Secondary extinctions") + xlab("Primary extinctions") + theme_bw()
R> g <- g + geom_point(data = Hypothesis$sims,
            aes(y = AccSecExt), color = "black") +
  geom_line(data = Hypothesis$sims,
            aes(y = AccSecExt, color = "black"), color = "black", size = 1.2)
R> g <- g + geom_point(data = Addmodel, aes(y = AccSecExt), color = "orange") +
  geom_line(data = Addmodel,
            aes(y = AccSecExt, color = "orange"), color = "orange", size = 1.2)
R> g
```
